# Improving cassava bacterial blight resistance by editing the epigenome

**DOI:** 10.1101/2022.01.04.474801

**Authors:** Kira M. Veley, Kiona Elliott, Greg Jensen, Zhenhui Zhong, Suhua Feng, Marisa Yoder, Kerrigan B. Gilbert, Jeffrey C. Berry, Zuh-Jyh Daniel Lin, Basudev Ghoshal, Javier Gallego-Bartolomé, Joanna Norton, Sharon Motomura-Wages, James C. Carrington, Steven E. Jacobsen, Rebecca S. Bart

## Abstract

Pathogens rely on expression of host susceptibility (S) genes to promote infection and disease. As DNA methylation is an epigenetic modification that affects gene expression, blocking access to S genes through targeted methylation could increase disease resistance. *Xanthomonas phaseoli* pv. *manihotis*, the causal agent of cassava bacterial blight (CBB), uses transcription activator-like20 (TAL20) to induce expression of the S gene *MeSWEET10a*. In this work, we direct methylation to the TAL20 effector binding element within the *MeSWEET10a* promoter using a synthetic zinc-finger DNA binding domain fused to a component of the RNA-directed DNA methylation pathway. We demonstrate that this methylation prevents TAL20 binding, blocks transcriptional activation of *MeSWEET10a in vivo* and that these plants display decreased CBB symptoms while maintaining normal growth and development. This work therefore presents an epigenome editing approach useful for crop improvement.

## Introduction

Susceptibility (S) genes present a liability within a host genome, as their exploitation is a common mechanism used by diverse classes of pathogens^1^. However, mutating these loci may have negative consequences as these genes are often necessary for normal plant growth and reproductive development^2–5^. In many cases, pathogens actively upregulate expression of S genes during infection. Plant pathogenic bacteria within the *Xanthomonas* and *Ralstonia* genera express transcription activator-like (TAL) effectors that activate transcription of S genes by recognizing and binding specific, largely predictable, promoter sequences (effector binding elements, EBEs) within the host genome^4,6–9^. Previous research demonstrated that gene editing can be used to prevent the exploitation of S genes by pathogens and that this results in increased resistance^10–14^. Epigenomic modifications can also influence gene expression. Because TAL binding can be inhibited by DNA methylation^15,16^, we hypothesized that targeting methylation to a TAL effector binding site would block induction of the S gene and result in increased resistance as measured by decreased disease symptoms.

In eukaryotes, cytosine DNA methylation is an important mechanism of epigenetic gene regulation^17^. In plants, the RNA-directed DNA methylation (RdDM) pathway is responsible for establishing *de novo* 5-methylcytosine methylation. Previous targeted methylation efforts leveraged naturally occurring epialleles from Arabidopsis that affect flowering time^18,19^. These studies used CRISPR-Cas9-based tools combined with components of the methylation pathways, either the catalytic domain of the plant methyltransferase DOMAINS REARRANGED METHYLTRANSFERASE 2 (DRM2) involved in RdDM or the bacterial CG-methyltransferase MQ1^20,21^. In addition, a number of RdDM proteins are effective in targeting methylation when fused with artificial zinc-fingers (ZFs), including DRM2 and DEFECTIVE IN MERISTEM SILENCING 3 (DMS3), a protein that in Arabidopsis targets RNA polymerase V binding to chromatin to trigger RdDM^22,23^. While CRISPR-based tools are highly specific, ZF-based targeting systems are preferable for some applications, as they generally require smaller transgenes and are not constrained by the location of PAM sites.

Cassava bacterial blight (CBB) is caused by *Xanthomonas phaseoli* pv. *manihotis*^24^ (*Xam*, aka. *Xanthomonas axonopodis* pv. *manihotis)* and occurs in all regions where cassava (*Manihot esculenta* Crantz) is grown^25,26^. Recently, a resistance gene was described that may provide some tolerance to certain strains of Xam^27^, however, currently, there are no strong broad spectrum resistance genes available to breeders. Symptoms of an infection begin with dark green, angular, “water-soaked” lesions on leaves (Fig. 1A) after which *Xam* moves through the vasculature causing leaf wilting and eventual death of the plant. *Xam* secretes 20 - 30 effectors, including multiple TAL effectors, into cassava cells^28^. We previously demonstrated that one of these TAL effectors, TAL20, is a major virulence determinant for this pathogen. TAL20 induces ectopic S gene expression of a SWEET (<u>S</u>ugars <u>W</u>ill <u>E</u>ventually be <u>E</u>xported <u>T</u>ransporters) family sugar transporter, *MeSWEET10a*^6,29^. Expression of *MeSWEET10a* during infection is required for water-soaking phenotypes and full virulence^6^. In wild-type cassava, the promoter of *MeSWEET10a* is not methylated^30^. Thus, we hypothesized that targeted methylation at the TAL20 EBE would prevent ectopic expression of *MeSWEET10a* and provide resistance to CBB (Fig. 1B).

**Fig. 1.**
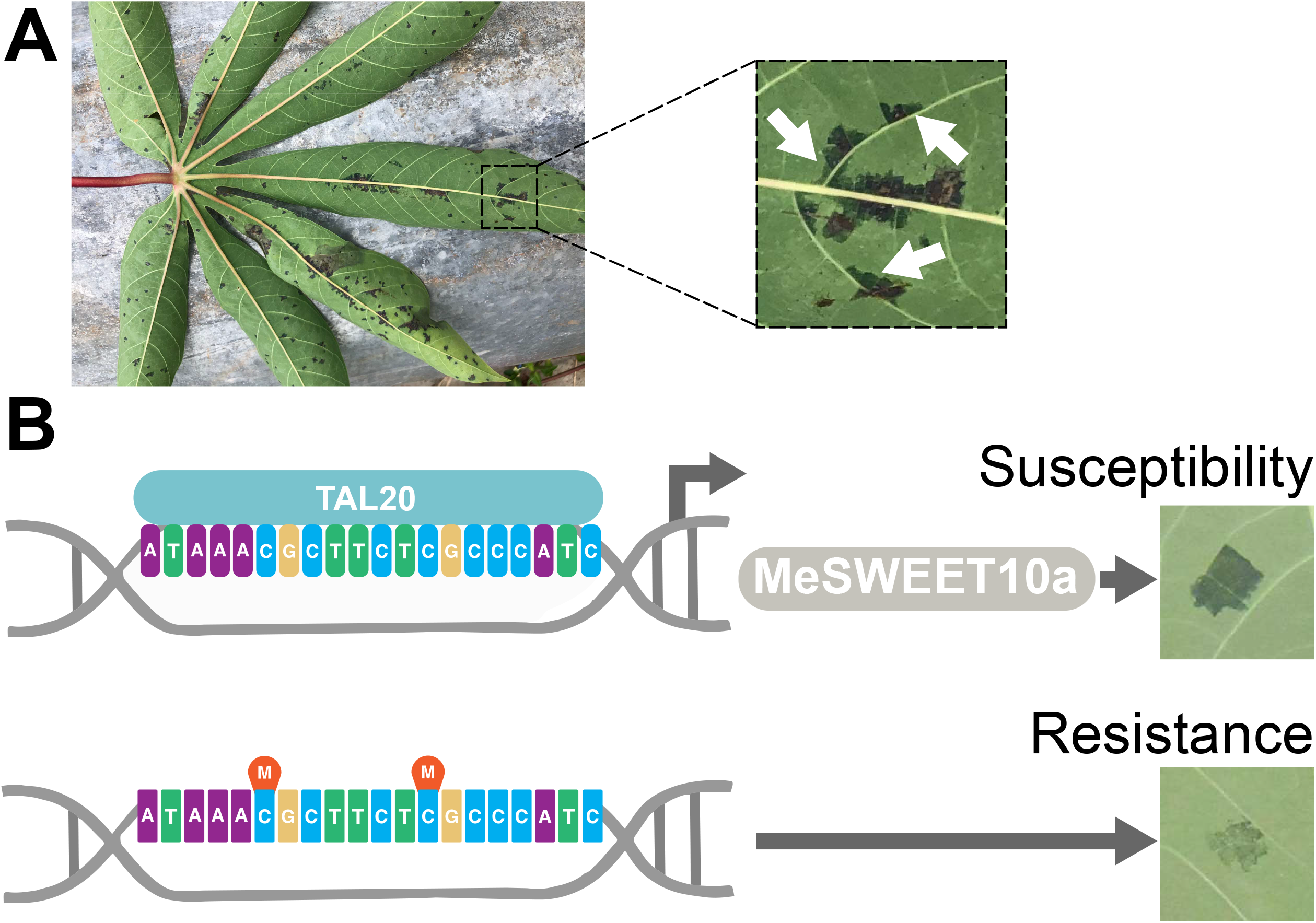
Overview of strategy to improve disease resistance using epigenetic modification. (**A**) Left: Example of CBB-infected cassava in a field (Uganda, 2019). Right: selected region from leaf on the left. Water-soaked regions are indicated with white arrows. (**B**) Graphical representation of epigenetic CBB disease resistance strategy. Top: In WT plants, TAL20 from *Xam* binds a specific sequence (EBE) present in the promoter of the S gene *MeSWEET10a*. Upon binding, TAL20 induces the ectopic expression of MeSWEET10a, a sugar transporter, which is required for establishment of disease. A typical ‘water-soaked’ lesion is shown to the right, an early indicator of CBB. Bottom: DNA methylation prevents TAL20 from binding the EBE. *MeSWEET10a* expression is not induced, and disease symptoms are reduced.

Here, we present epigenetic editing as a useful approach toward crop improvement. We show directed DNA methylation to the TAL20 EBE within the *MeSWEET10a* promoter using a ZF fused to DMS3. We demonstrate that cytosine methylation inhibits TAL20 binding to the EBE, preventing transcriptional activation of *MeSWEET10a* and decreasing bacterial blight symptoms in cassava plants.

## Results

### Establishing methylation at the *MeSWEET10a* promoter

We first tested the effect of methylation on TAL20 binding to the EBE *in vitro* using an Electrophoretic Mobility Shift Assay (EMSA) (Supplementary Fig. 1). As shown previously^6^, purified 6xHis-TAL20_Xam668_ binds the unmethylated EBE oligo (lane 2) and, as expected, providing an excess of unlabeled, unmethylated oligo outcompeted for binding (lane 5). Conversely, 6xHis-TAL20_Xam668_ does not efficiently bind the methylated oligo (lane 3), nor did the methylated oligo compete for TAL20 binding, even when supplied in excess (lanes 6 and 7). Therefore, the EBE of *MeSWEET10a* was deemed a good candidate for engineering epigenetic disease resistance. Of the RdDM factors tested in *Arabidopsis thaliana*, DMS3 was shown to be the most effective at delivering methylation to a given locus^23^. Transgenic cassava plants were generated expressing an artificial ZF targeting the EBE of *MeSWEET10a* fused to Arabidopsis DMS3 (Fig. 2A, Supplementary Figs. 2-7). The targeted region of the *MeSWEET10a* promoter was assessed for methylation using amplicon-based bisulfite sequencing (ampBS-Seq) (Fig. 2B, Supplementary Figs. 2-7). Plants expressing the ZF alone showed no methylation at or around the EBE of *MeSWEET10a*. In contrast, all plants expressing DMS3-ZF contained high levels of ectopic CpG methylation within and near the ZF target, including the only two CpG sites within the EBE (Fig. 2B, Supplementary Figs. 2-7). The specific analysis that the plants were used for in each Supplementary Fig. 2-7 are indicated in the title of the figure. Plant phenotypes were evaluated under greenhouse and field conditions using our standard methods (see methods for details). These analyses indicated that methylation did not grossly affect root and shoot growth and development (Fig. 2C, D, Supplementary Fig. 8). Whole Genome Bisulfite Sequencing (WGBS) was used to assess specificity of the DMS3-ZF. High levels of *de novo* DNA methylation were observed within the target region, similar to the ampBS-seq result (Fig. 2B, Fig. 3A), while DNA methylation within an extended region of 5 kb on either side was comparable between the DMS3-ZF-expressing lines and the ZF alone control (Fig. 3A, B). Additionally, CG, CHG, and CHH methylation across all chromosomes, as well as within genes, was similar between the DMS3-ZF-expressing lines and ZF alone control (Fig. 3C, D, Supplementary Fig. 9, and Supplementary Table 1). We investigated potential off-targets by predicting ZF binding sites with up to 4 mismatches within the target sequence (5’-CGCTTCTCGCCCATCCAT - 3’). This yielded 1,542 potential off-target regions genome-wide (Supplementary Fig. 10A). We compared methylation levels of these potential off-targets, plus 100bp flanking sequences, in WT and the two DMS3-ZF-expressing lines 133 and 204. No significant differences were detected (Supplementary Fig. 10B). As a complementary approach to identify potential non-specific activity, differentially methylated regions (DMRs) (CG variation > 0.4, CHG variation > 0.2, and CHH variation > 0.1, respectively) were obtained by comparing the two DMS3-ZF lines with the WT control. This method successfully detected two hypermethylated DMRs in the targeted MeSWEET10a region (Fig. 3A). However, most of the other detected DMRs were hypoDMRs (i. e. regions showing decreased methylation in the DMS3-ZF lines) rather than hyperDMRs (increased methylation in the DMS3-ZF lines). An overall decrease in genome methylation has been attributed to the plant tissue culture process in cassava^31^ and in many other plant species including maize^32^, rice^33^ and oil palm^34^. Cross referencing the observed DMRs with the 1,542 potential off-targets revealed that very few of DMRs (particularly hyperDMRs) fell into off-target regions, and the chance was indistinguishable from the random control regions (Supplementary Fig. 10C, Supplementary Table 2). Together, these results demonstrate a reasonable level of specificity for targeted DMS3-ZF-mediated methylation in cassava.

**Fig. 2.**
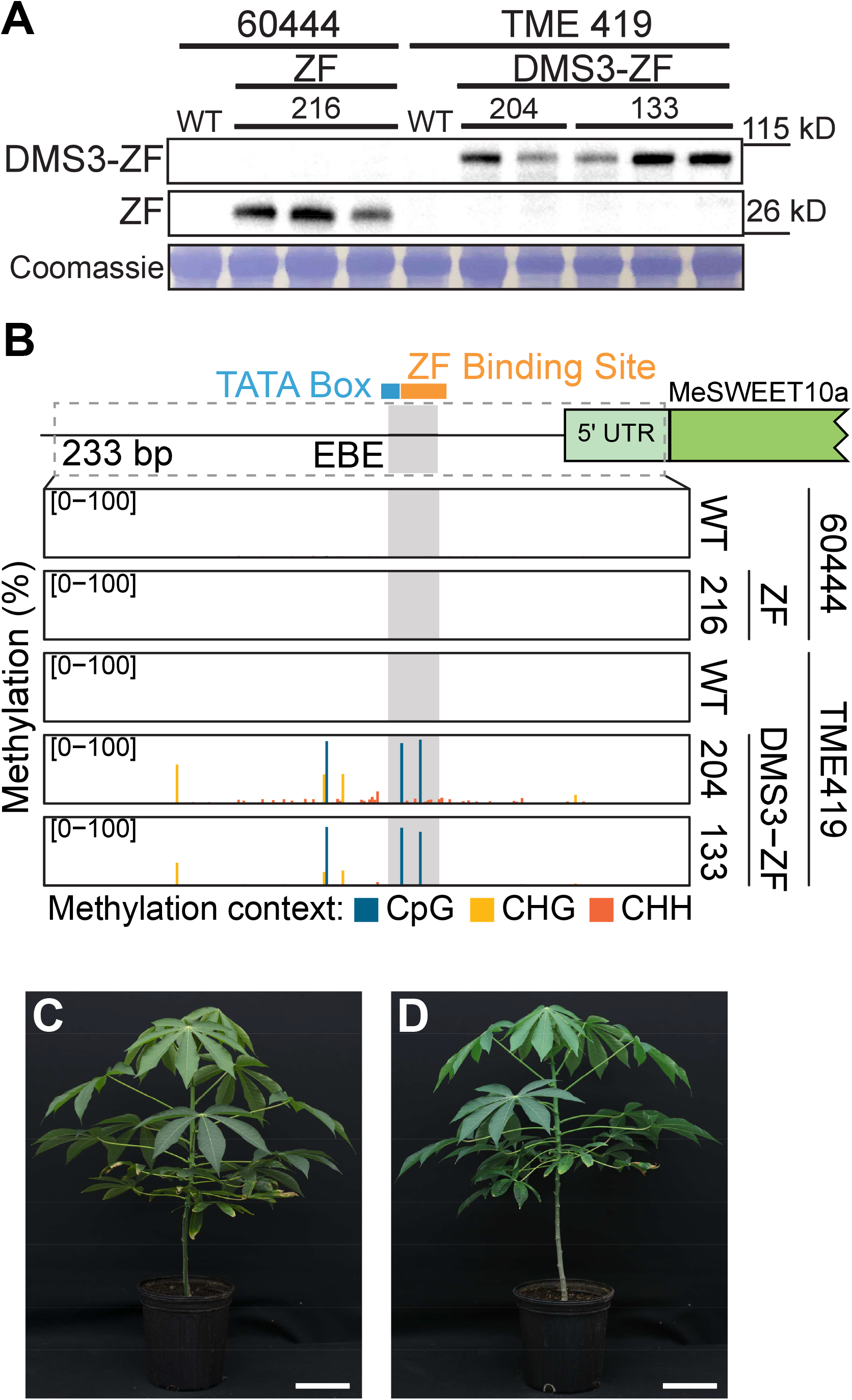
DMS3-ZF expression results in CpG methylation at the EBE *in vivo*. (**A**) Expression of transgenes in individual plants from two independent DMS3-expressing transgenic lines (133 and 204) as well as a ZF-only negative control line (216). Cassava variety information for each sample is given above the lanes. First two rows: representative western blots (anti-FLAG) showing expression of the ZF (ZF-3xFLAG) protein with (top) and without (middle) DMS3. Relevant size standards are shown to the right (kD). Bottom: Coomassie Brilliant Blue stained Rubisco, loading control. Uncropped images of the western and Coomassie are shown in **Supplementary Fig. 5A**. (**B**) Representative PCR-based bisulfite sequencing (ampBS-seq) results from samples shown in A. Details results for these samples are shown in **Supplementary Fig. 5A**. Top: Graphical depiction of *MeSWEET10a* promoter region assessed for methylation. The EBE (grey), a presumed TATA box (blue) and the ZF binding site (orange) are indicated. The predicted 5’ UTR and *MeSWEET10a* transcriptional start site are shown in green. The area within the dotted lined box (233 bp) was subjected to ampBS-seq. Bottom: CpG, CHG, and CHH DNA methylation levels (percent, y-axis) of the *MeSWEET10a* promoter (EBE, grey) measured by ampBS-seq with and without DMS3-ZF. Background of tissue for each plot is indicated to the right. Methylation was called for cytosines with greater than 500 reads mapped. The results in panels A and B are representative examples of the six independent experiments presented in **Supplementary Figs. 2-7**. (C) Representative wild-type (TME419) and (**D**) DMS3-ZF-expressing (line #133) plants. Scale bar = 14 cm. Additional data quantifying growth in the field are shown in **Supplementary Fig. 8**.

**Fig. 3.**
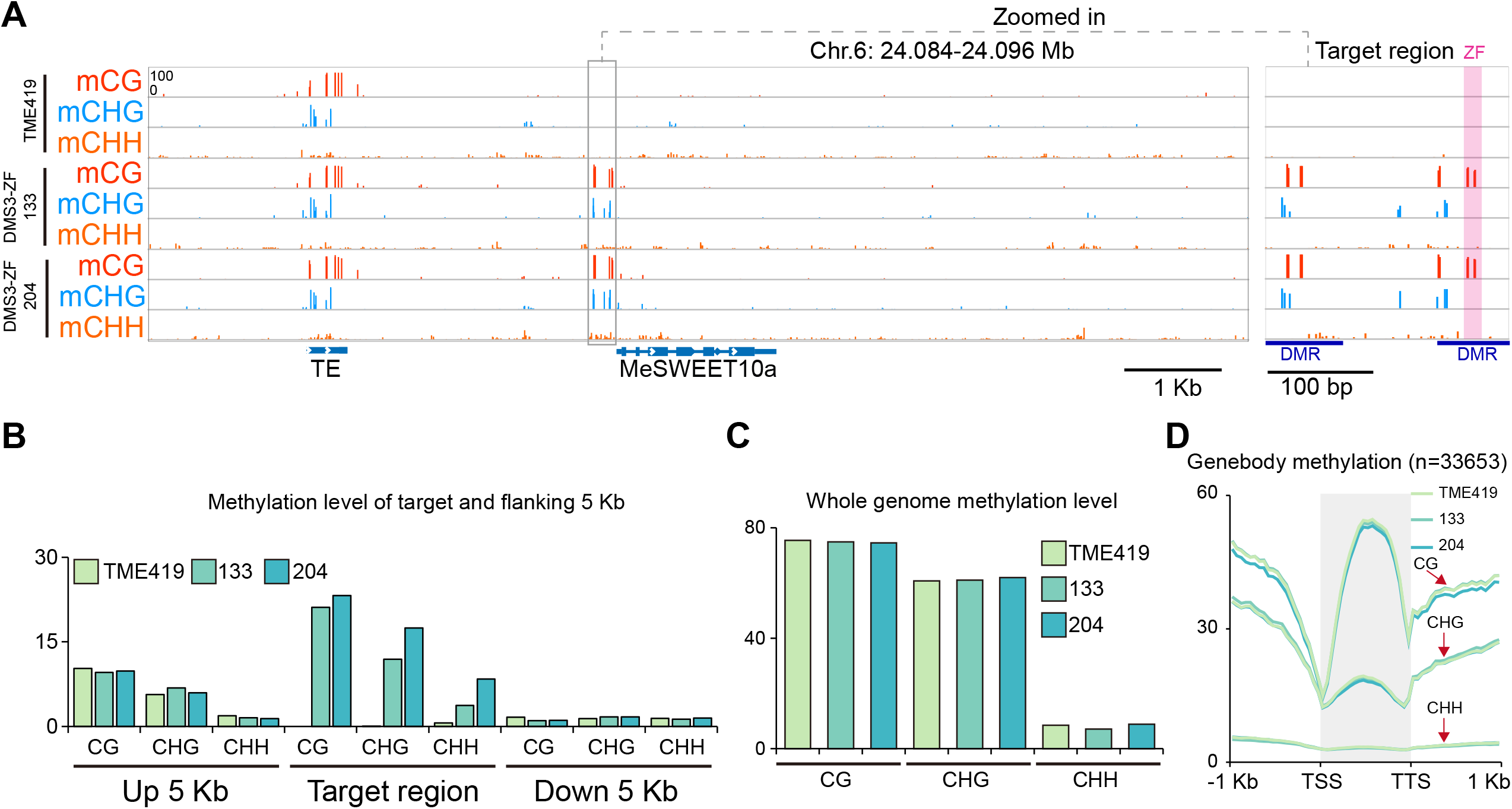
Targeted methylation at MeSWEET10a shows high specificity. **(A)** Distribution of CpG, CHG, and CHH methylation over the targeted and flanking 5 kb regions of the *MeSWEET10a* promoter in wild-type TME419 and two independent DMS3-ZF lines (transgenic lines 133 and 204). Grey box indicates the targeted region within the *MeSWEET10a* promoter. ZF binding site is shaded within the pink box. Two detected hypermethylated differentially methylated regions (DMR) are also indicated. **(B)** Summary of CpG, CHG, and CHH methylation level of targeted and flanking 5 kb regions of the *MeSWEET10a* promoter in wild-type TME419 and the two independent DMS3-ZF lines. **(C)** Whole genome CpG, CHG, and CHH methylation level of wild-type TME419 and the two representative DMS3-ZF lines. **(D)** Metaplot of CpG, CHG, and CHH methylation over the gene of protein coding genes (n = 33,653) and flanking 1 kb regions in wild-type TME419 and the two independent DMS3-ZF lines. Transcriptional start sites (TSS) and transcriptional termination sites (TTS) are indicated.

### Cytosine methylation prevents *MeSWEET10a* transcriptional activation by TAL20 *in vivo*

We next tested the hypothesis that CpG methylation at the TAL20 EBE would prevent activation of *MeSWEET10a in vivo*. Compared to controls, plants from independent DMS3-ZF-expressing lines showed a significant lack of *MeSWEET10a* induction in response to *Xam* infection (Fig. 4A). The suppression of *MeSWEET10a* induction occurred in independently derived transgenic lines and across independent experiments. This suppression did not correlate with increased expression of DMS3-ZF (Supplementary Fig. 11). These data indicate that methylation prevents TAL20-mediated activation of *MeSWEET10* expression.

**Fig. 4.**
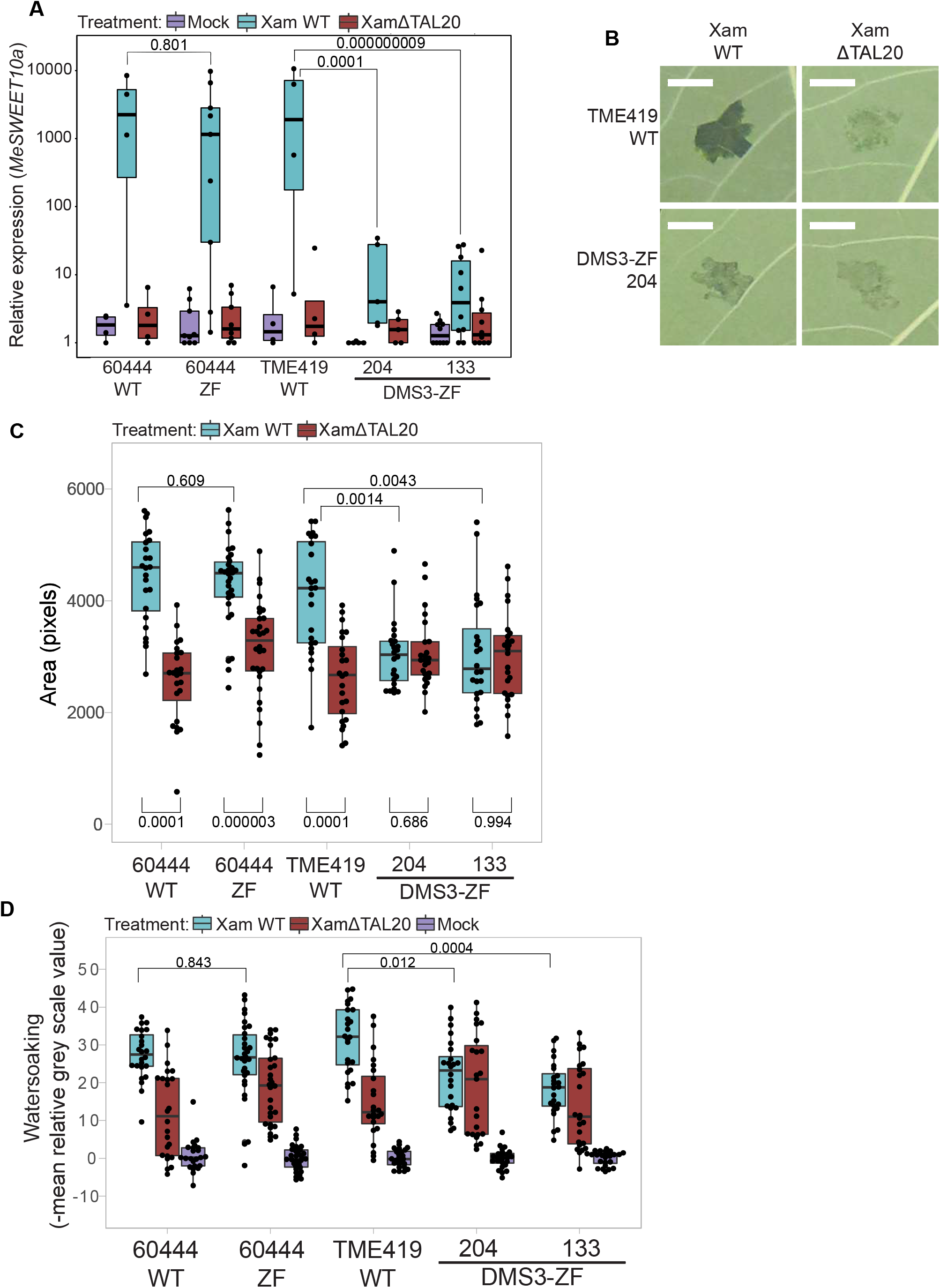
Effect of methylation on CBB disease phenotypes in cassava. (**A**) Normalized *MeSWEET10a* expression (y-axis, log10 scale) in WT and transgenic lines as determined by RT-qPCR. The cassava genes GTPb (Manes.09G086600) and PP2A4 (Manes.09G039900) were used as internal controls. Boxes are colored according to Xanthomonas treatment. Biological replicates (black dots) included in each background (x-axis) are as follows: n = 4, 9, 4, 5, 10 examined over 4 independent experiments. The n included in each treatment group for each background are consistent. Horizontal black line within boxes indicates the value of the median while the box limits indicate the 25th and 75th percentiles as determined by R software; whiskers extend 1.5 times the interquartile range (1.5xIQR) from the 25th and 75th percentiles. Two-sided Welch’s *t*-test *p-*values are noted above brackets within plot. (**B**) Representative images of water-soaking phenotype of leaves from TME419 WT and DMS3-ZF-expressing plants. Images were taken 4 days post-infection with either *Xam668* (XamWT) or a *Xam668* TAL20 deletion mutant (Xam_Δ_TAL20). Scale bar = 0.5 cm. (**C**) Observed area (pixels, y-axis) of water-soaking from images of Xam-infiltrated leaves (backgrounds, x-axis) 4 days post-infiltration. Images from (**B**) and **Supplementary Fig. 11** are included in dataset. Calculated *p-*values (Kolmogorov-Smirnov test) are shown above brackets within plot. (**D**) Intensity of water-soaking phenotype (y-axis) of region measured in panel **C**. The negative mean grey-scale value for the water-soaked region relative to the average of the mock-treated samples within the same leaf is reported (see methods for details). Calculated *p*-values (two-sided Kolmogorov-Smirnov test) are shown above brackets within plot. Both box plots: Biological replicates (black dots) included in each background (x-axis) are as follows: n = 24, 30, 24, 24, 24 examined over three independent experiments.Horizontal black line within boxes indicates the value of the median while the box limits indicate the 25th and 75th percentiles as determined by R software; whiskers extend 1.5 times the interquartile range (1.5xIQR) from the 25th and 75th percentiles.

### Epigenomic editing at *MeSWEET10a* decreases cassava bacterial blight symptoms

To quantify the effect of methylation at the *MeSWEET10a* EBE on CBB disease *in vivo*, we measured “water-soaked” lesions (Fig. 1A, Fig. 4B, Supplementary Fig. 12), which are the most obvious early disease symptom and requires induction of *MeSWEET10a*^6,35^. Quantitative analysis of images from *Xam*-infiltrated leaves revealed that plants with methylation at the EBE had a reduced infected area compared to wild-type (Fig. 4C). Further, the area of infection in DMS3-ZF leaves infiltrated with wild-type *Xam* was comparable to leaves infected with *Xam* lacking TAL20 (Xam_Δ_TAL20). No significant difference in bacterial population was observed in the DMS3-ZF lines (Supplementary Fig. 13). In addition to a decrease in lesion size, the intensity of water-soaking at and around the infiltration site was also reduced in EBE methylated plants. Grey-scale values from images of lesions were compared to mock-inoculated sites on the same leaf, where more water-soaking equates to a darker appearance. Water-soaking intensity was reduced in plants with methylation at the EBE (Fig. 4D). These results demonstrate that methylation of the EBE site within the promoter of the S gene *MeSWEET10a* results in a reduction in CBB-related symptoms.

## Discussion

In summary, we show that DMS3 paired with an artificial ZF caused targeted, *de novo* establishment of methylation at the EBE of *MeSWEET10a*, blocking TAL20-induced ectopic expression of *MeSWEET10a* and resulted in decreased CBB disease symptoms. Cassava is typically clonally propagated, and we have monitored methylation over time throughout the course of this study (three-four years maintenance in tissue culture and in the greenhouse). We conclude that the methylation is stable in the presence of the transgene through clonal propagation. Because the components of the technology presented here do not derive from cassava (Arabidopsis DMS3 paired with a synthetic zinc finger), it is reasonable to hypothesize that this approach may be similarly successful in other crop species. Gene editing represents an existing, powerful strategy that can achieve similar outcomes to the work presented here, and we expect this will continue to be the favored approach for many applications^5,10–14,36–40^. At the same time, as targeted methylation technologies continue to mature, there will likely be cases where it may be preferred. For example, DNA methylation may be applied as a gene expression dial, opposed to an on/off switch. Similarly, as DNA methylation does not change the DNA sequence, it may be possible to use DNA methylation changes to achieve expression patterns in a tissue- or environment-specific manner. This might be achieved in transgenic plants by driving the methylation reagents using a tissue- or environment-specific promoter. Additionally, a key near-term future goal will be to study the heritability of DNA methylation in the absence of the targeting transgene^41,42^. It seems likely that the size of the region with de novo methylation, the type of methylation (CG, CHG, CHH), and the specific location in the genome may all affect heritability. These factors will each need to be investigated, and while there is value in pursuing this work in the crop of interest, we expect that much can be learned using model systems. Ultimately, the practicality of this strategy will be determined by the heritability and stability of epigenetic edits, and this is likely to be application- and target-specific. Nonetheless, the transgenic epigenetic approach shown here may be useful in a variety of other crop improvement programs.

## Methods

### Construct design and cloning

The genomic coding sequence of Arabidopsis *DMS3* (AT3G49250), including promoter, was cloned into the pEG302 backbone as described^23,44^. The ZF targeting the EBE of *MeSWEET10a* was designed as previously described^22^ to target the sequence 5’-CGCTTCTCGCCCATCCAT-3’. CodeIT (zinc.genomecenter.ucdavis.edu:8080/Plone/codeit) was used to generate the amino acid in silico (linker type “normal”). The resulting amino acid sequence was plant codon optimized, synthesized by IDT and cloned at the 3’ end of *DMS3* (AT3G49250) in pEG302. A hygromycin resistance cassette was cloned into an available PmeI site for selection in cassava. Vector sequences and maps are available in supplementary materials.

### Production of plants and growth conditions

The transgenic cassava line expressing the ZF with no methylating factor was generated for a different project, and thus a different background was used: 60444 (ZFΔDMS3) versus TME419 (DMS3-ZF). The promoter region of interest is identical in both genetic backgrounds, as verified by Sanger sequencing (Extended Data Fig. 14). Transgenic cassava lines of cultivar TME419 and 60444 were generated and maintained *in vitro* as described previously^45^ with some modifications, as follows: transgenic FEC cells were selected for resistance using hygromycin (20 mg / ml) on spread plates (also containing 125 mg / L cefotaxime). On stage 1, 2, and 3 plates, 20 mg / ml hygromycin was used as well. Growth chamber settings used for tissue culture: 28°C +/- 1°C, 75 μmol·m^-2^·s^-1^ light; 16 hrs light / 8 hrs dark. *In vitro* plantlets were propagated and established in soil on a misting bench with domes. Once established, plants were removed from the misting bench and acclimated to greenhouse conditions (28°C; humidity >50%; 16 hrs light / 8 hrs dark; 1000 W light fixtures supplementing natural light levels below 400 W / m^2^). 1000 W light fixtures supplementing natural light levels below 400 W / m^2^). For field-grown plants, individuals were first established in the greenhouse (natural light and temperature, 17-28°C, humidity >50%) for 29 weeks then transitioned outside and planted 3 feet apart (field location 19.640, -155.083).

### PCR bisulfite sequencing (ampBS-seq) library preparation and analysis

PCR-based BS-seq (ampBS-seq) of the *MeSWEET10a* promoter was performed as described^21^. Leaf tissue and DNA extracted using the GenElute™ Plant Genomic DNA Miniprep Kit (Sigma) and sodium bisulfite treated using the Epitect Bisulfite Conversion kit (QIAGEN). The region of interest within the *MeSWEET10a* promoter was amplified using the primers in Extended Data Table 3, giving a 233bp product. Libraries were generated from purified PCR products amplified from the bisulfite-treated DNA, using an Ovation Ultralow V2 kit (NuGEN) or a Kapa DNA HyperPrep Kit (Roche) in combination with IDT UD barcodes (Illumina). iSeq 100 (Illumina) was used to sequence the libraries. Raw sequencing reads were aligned to the region of interest using the epigenomic analysis plugin for CLC Genomics Workbench 20.0 (QIAGEN). Reads shorter than 125 bp or longer than 152 bp were discarded. Reads were mapped to the following sequence: ATATTGTTTTTTTTTTTTTTAAAAAAAATAATAAAAGAAATAAGGTTACTGTTACATT GACATATTTTATTCACTTTAATCATGCATGCAACTTGACTTCATTCCGTTCCCTGGAT TCCTCCCCTATATAAACGCTTCTCGCCCATCCATCATTGCACAACATAGCTAGAGTTT CCTCTTGAGAAAGAGAGTTTCCTCTGCATAAGGGAAAGAGAGTTTTTATTATAGTYG GA. For mapping, the following parameters were used: non-directional, no masking, match score = 1, mismatch cost = 2, affine gap cost, insertion open cost = 6, insertion extend cost = 1, deletion open cost = 6, deletion extend cost = 1, length fraction = 0.9, similarity fraction = 0.9. Methylation was called for cytosines with greater than 500 reads mapped and the resulting methylation level was plotted using the provided script (MeSWEET10a_07272021-for_publication.0920.R).

### Whole Genome Bisulfite sequencing (WGBS) library preparation and analysis

Genomic DNA from samples were end-repaired and ligated with TruSeq LT DNA single adapters (Illumina) using Kapa DNA HyperPrep kit (Roche). Adapter-ligated DNA was converted with EpiTect Bisulfite Conversion Kit (Qiagen). Converted DNA was PCR-amplified by MyTaq polymerase (Bioline) for 12 cycles. Library quality and size were determined by D1000 ScreenTape (Agilent) and libraries were subsequently purified by AMPure XP beads (Beckman Coulter). Library concentrations were measured with Qubit dsDNA Broad-Range Assay kit (ThermoFisher). Libraries were sequenced on NovaSeq 6000 sequencer (Illumina). WGBS reads were mapped to the TME7 phase0 genome with BSMAP (v2.90) allowing 1 mismatch and 1 best hit (-v 1 -w 1)^46,47^. Duplicated reads were removed with SAMtools (v1.3.1)^48^. Reads with three or more consecutive methylated CHH sites were considered as unconverted reads and removed in the following analysis. Conversion rate was estimated by calculating methylation level of the chloroplast genome. DNA methylation level at each cytosine was calculated by the number of methylated C vs. total C and T account. Methylation levels of each chromosome were determined by calculating CpG, CHG, and CHH level of 100 kb bins. Differentially Methylated Regions (DMRs) were called using BSMAP (v2.90) with p < 0.01, where the differences in CG, CHG, and CHH methylation were at least 0.4, 0.2, and 0.1, respectively. Methylation track files were visualized with Integrative Genomics Viewer (IGV, v3.0).

### Off-target analysis

ZF off-target sites were predicted by Cas-OFFinder by searching cassava genome with target site (5’-CGCTTCTCGCCCATCCAT-3’) allowing 4 mismatches and no PAM site^49^. Methylation levels of off-target sites and 100 bp flanking sequences (50 bp upstream and 50 bp downstream) were compared between samples. Sequence consensus analysis of potential off-target sites was performed with WebLogo (v3.6.0)^50^.

### SDS-PAGE gel and western blotting

All individuals used for qPCR and disease assay experiments were checked for construct expression (Extended Data Figs. 3 (qPCR1), 4 (qPCR2), 5 (Water-soaking), 6 (qPCR3), 7 (Water-soaking, qPCR4), 8 (Water-soaking)). Tissue (100 mg) was collected from each individual plant to be used in an experiment in a 1.5 ml microcentrifuge tube and frozen in liquid nitrogen along with four 3mm glass beads. A tissue lyser (TissueLyser II, QIAGEN) was used to homogenize the sample (30/second frequency, 2 minutes, performed twice with sample rotation between runs). 400 ul of 1x Laemmli Sample buffer (.0625M Tris-HCl pH 6.8, 2% SDS, 10% glycerol, 100 mM DTT) was added to each sample, then samples were boiled 90-100°C for 4 min, centrifuged for 2 min and then either run immediately on a gel or frozen for storage at - 80°C. For SDS-PAGE, 10ul of each sample was run on a 12% TEO-Tricine Precast Gel (RunBlue™, AbCam, ab119203) then transferred to PVDF membrane (ThermoFisher Scientific). For western blotting, monoclonal ANTI-FLAG® M2-Peroxidase (HRP) antibody (A8592, Sigma) was used (1:2000 dilution in 5% milk in TBST). HRP activity was assessed using SuperSignal™ West Femto substrate (ThermoFisher Scientific). Chemiluminescence was detected using the ChemiDoc XRS+ System equipped with Image Lab 6.1 software (Bio-Rad). The molecular weight of protein bands was estimated by comparison with the BenchMark™ Pre-stained Protein Ladder (Thermo Fisher Scientific). The abundance of the 3xFLAG-ZF proteins were quantified from the band intensities also using Image Lab Software. Coomassie staining was performed on membranes using Coomassie brilliant blue (0.025% (w/v) Coomassie brilliant blue R-250 (ThermoFisher Scientific) in 40% methanol/7% acetic acid (v/v)). Staining was performed for 3 min followed by destain (solution: 50% methanol/7% acetic acid (v/v)) for 30 min. Then membranes were rinsed with water and allowed to dry overnight before imaging.

### Electrophoretic mobility shift assay (EMSA)

For protein purification, *TAL20*_*Xam668*_ was cloned into pDEST17 (Thermo Fisher Scientific) using Gateway™ technology, resulting in the addition of a 6xHis-tag at the N-terminus of the protein upon expression. 6xHis-TAL20_Xam668_ was expressed and purified from *E. coli* BL21 cells using HisPur™ Ni-NTA Spin Columns (Thermo Fisher Scientific). Protein size and purity was assessed by SDS-PAGE (4-12% gradient gel) followed by western blot (Extended Data Fig. 1A) using an anti-His antibody (His-Tag (27E8) Mouse mAb (HRP Conjugate), Catalog #9991, Cell Signaling Technology). For the EMSA, the sequence previously shown to be able to be bound by TAL20_Xam668_ ^6^, with and without methylation, was synthesized (Integrated DNA Technologies) and biotin-labeled (Pierce™ Biotin 3’ End DNA Labeling Kit, Thermo Fisher Scientific). The sequence used is as follows, with methylated cytosines indicated by underlining: sense 5’-CCCTATATAAA<u>C</u>G<u>C</u>TT<u>C</u>T<u>C</u>G<u>CCC</u>AT<u>C</u>CATCATT-3’, anti-sense: 3’-GGGATATATTTG<u>C</u>GAAGAG<u>C</u>GGGTAGGTAGTAA-5’. EMSA was performed with the Light Shift® Chemiluminescent EMSA Kit (Thermo Fisher Scientific) according to the manufacturer’s protocol. Biotin-labeled oligonucleotides were mixed with 0, 20, 200, 300 and 1000 fmol of purified TAL20_Xam668_ effector protein as indicated in the chart in Extended Data Fig. 1B. All binding reactions were incubated at room temperature for 20 min. For competition assays, unlabeled oligonucleotides were pre-incubated with other reaction components for an additional 20 min. Gel electrophoresis was performed on a 6% native polyacrylamide gel.

### Bacterial inoculations

*Xanthomonas* strains were grown on plates containing necessary antibiotics for 2-3 days at 30°C. The *Xanthomonas* strains used and their antibiotic resistances are as follows: 1. *Xam668* (rifampicin 50 μg / ml) and 2. *Xam668*_Δ_*TAL20* (suicide vector knockout^6^, tetracycline 5 μg / ml, rifampicin 50 μg / ml). Bacteria were scraped from the plates into 10 mM MgCl_2_ to either OD_600_ = 1.0 (RT-qPCR) or OD_600_ = 0.01 (water-soaking) concentrations according to established lab protocols. Leaves (2-3 weeks after plants were transferred to a greenhouse) were inoculated with a 1.0 ml needleless syringe using one bacterial strain per lobe, three injection points on each of 2-3 leaves. After inoculation, plants were kept under fluorescent light (12 hr day / night light cycle) at 27°C for either 48 hours (RT-qPCR and ampBS-seq) or 4 days (water-soaking).

### RNA extraction and RT–qPCR analysis

For RNA, 48 hours post-infiltration, samples were taken using an 8 mm cork borer at each infiltration site, frozen in liquid nitrogen, ground to a fine powder (3 mm glass beads added to tissue, ground using tissue lyser (TissueLyser II, QIAGEN), 30/second frequency, 2 minutes, performed twice with sample rotation between runs) and total RNA was extracted (Spectrum™ Plant Total RNA Kit, Sigma). Technical replicates were pooled, thus each RNA sample contained three injection sites from one leaf. 1.0 μg DNase-treated RNA was reverse-transcribed to cDNA using SuperScript™ III Reverse Transcriptase (Thermo Fisher Scientific) according to the manufacturer’s instructions.

Quantitative reverse transcription PCR (RT-qPCR) was done with SYBR Select Master Mix (Thermo Fisher Scientific) on a CFX384 Touch Real-Time PCR Detection System (Bio-Rad). Primers specific for cassava GTPb (Manes.09G086600) and PP2A4 (Manes.09G039900) genes were used as the internal controls to calculate relative expression levels (Table S1)^51,52^. Normalized relative expression was calculated using formulas described^53^.

### Water-soaking area and intensity quantification, image analysis

Cassava leaves were infiltrated with either 10 mM MgCl_2_ alone (mock) or containing *Xanthomonas* (*Xam668* strains with and without TAL20) as described above and imaged using a Canon EOS Rebel T5i camera with a 15-85mm lens at 0- and 4-days post-inoculation. Images were grey balanced using a X-Rite Passport by estimating the saturation of the six grey chips and lowering the brightness accordingly. Gray corrected images were uploaded to FIJI^54^ and duplicated. In order to define the infected area, visible water-soaking spots were manually outlined on the duplicate image using the pencil tool (color: #ff00b6 and size 2). The outlined images were converted from RGB to LAB and split to obtain the A color channel. The A channel images were thresholded, converted to a mask and the mask for each spot was added to the ROI manager using the analyze particle tool. The ROI masks were applied to the original RGB grey corrected images. In order to define the background color of a given leaf, mock-infiltrated spots (no water-soaking) were added to the ROI manager using an arbitrarily sized rectangle selection tool consistently set to a W = 51 and H = 61. Area and grey scale mean data were obtained for each infiltrated spot using the FIJI measure tool. Grey balancing using the X-Rite Passport is a coarse adjustment and finer standardization is performed post image processing using statistical methods. Grey value standardization was achieved by estimating the grand mean of all grey values in each image and centering those values to the grand mean of all images by using model residuals. Further, the residuals within each image were re-centered with respect to the mock treatment by subtracting the mean grey value of mock from all other features.

### Bacterial growth assay

Cassava leaves were infiltrated with either 10 mM MgCl_2_ alone (mock) or containing *Xanthomonas* (*Xam668* strains with and without TAL20) as described above. Leaf punch samples were taken at the site of infiltration using a 5 mm cork borer at 0- and 4-days post-inoculation. For day-0 samples, infiltrated spots were allowed to dry down prior to processing. Individual leaf punches were transferred to 2 mL Eppendorf Safelock tubes with 200 ul of 10 mM MgCl_2_ and three disposable 3 mm Propper solid glass beads. Samples were ground with a Qiagen Tissuelyzer at 28 hZ for 3 minutes. 200 ul of the ground sample were transferred to the first column of a labelled 96-well plate. Serial dilutions were performed by transferring 20 ul of the non-dilute sample (10^1^) to the next well containing 180 ul of 10 mM MgCl_2_. Samples were serially diluted to 10^4^ for day-0 and 10^6^ for day-4. 10 ul of each serial dilution was spread onto labelled quadrants of a NYG plate with cycloheximide and the appropriate antibiotics for the infiltrated bacterial strain. Plates were incubated at 30°C for 2-3 days, and the number of colonies were counted. Colony forming Units (CFU) reported in this manuscript were transformed by sample area (CFU/cm^2^ where cm^2^ = 0.52).

### Storage root production and measurements

Storage root production and analysis was conducted with some modification as described^43^. Briefly, plants were maintained in small, 2-inch pots for 11 weeks in order to induce storage root production. The same greenhouse conditions were used as described above. After 11 weeks, roots were removed from pots, washed to remove soil, and storage roots isolated from the root mass and weighed for fresh weight measurements. For starch content estimation (approximately 80% of storage root weight), the storage roots were peeled, cut into approximately 2 cm cubes, placed into a 50 ml conical tube, and frozen in dry ice and stored at - 80°C prior to lyophilization. Samples were lyophilized for two days, then weighed to obtain dry weight.

## Data availability

All data and constructs are available on request. All amplicon-based bisulfite sequencing data, western blots, and images used for imaged-based analysis of water-soaking disease symptoms are available in a FigShare repository (DOI: 10.6084/m9.figshare.16887934, https://figshare.com/s/9b2cf4f3cdf894e67baf). Whole genome bisulfite sequencing (WGBS) data is available at GEO (https://www.ncbi.nlm.nih.gov/geo/query/acc.cgi?acc=GSE187022): GSM5667182 (WGBS data of wild type TME419), GSM5667183 (WGBS data of DMS3-ZF line 133), GSM5667184 (WGBS data of DMS3-ZF line 204). Source data are provided with this paper.

## Code availability

R code for generating plots are available in a FigShare repository (DOI:10.6084/m9.figshare.16887934, https://figshare.com/s/9b2cf4f3cdf894e67baf).

## Acknowledgments

We are grateful to our scientific advisory board, including Drs. Morag Ferguson, Nathan Springer, and Nigel Taylor. We also acknowledge Dr. Noah Fahlgren, Director of the Data Science lab at DDPSC, and the additional members of our labs who contributed to this work in various ways, especially Ke Ke, Molly Kuhs, Joshua Sumner, and Dr. Xinggou Zheng. We also thank David Segal for assistance with the design of the artificial zinc finger. This work was funded by the Bill and Melinda Gates Foundation (OPP1125410, RBS, SEJ, JCC) and the National Science Foundation (GRFP DGE-2139839 and DGE-1745038, KE).

## Author contributions

Conceptualization: SMW, JCC, SEJ, RSB

Methodology: KMV, KE, GJ, ZZ, SF, MY, KBG, JCB, ZJDL, BG, JGB, JCC, SEJ, RSB

Investigation: KMV, KE, GJ, ZZ, SF, MY, KBG, JCB, ZJDL, BG, JGB, JN

Visualization: KMV, KE, ZZ, MY, KBG, JCB

Funding acquisition: SMW, JCC, SEJ, RSB

Project administration: KBG, JN, SMW, JCC, SEJ, RSB

Supervision: SMW, JCC, SEJ, RSB

Writing – original draft: KMV, ZZ, RSB

Writing – review & editing: KMV, KE, GJ, ZZ, SF, MY, KBG, JCB, ZJDL, BG, JGB, JN, SMW, JCC, SEJ, RSB

## Competing interests

Authors declare that they have no competing interests.

## Notes

### Competing Interest Statement

The authors have declared no competing interest.

### Summary of Updates

This article has been updated to include more off-target analysis in addition to other, more minor, changes.

