## Supplemental figures and tables for "Improving cassava bacterial blight resistance by editing the epigenome"

**This file includes:**

Supplementary Figs. 1-14

Supplementary Tables 1-3

**Other supplementary information this manuscript includes the following:**

All amplicon-based bisulfite sequencing (ampBS-seq) data, western blots, images used for imaged-based analysis of water-soaking disease symptoms, and R code for generating plots are available in a FigShare repository (DOI: 10.6084/m9.figshare.16887934, <https://figshare.com/s/9b2cf4f3cdf894e67baf>). Whole genome bisulfite sequencing (WGBS) data is available at GEO (<https://www.ncbi.nlm.nih.gov/geo/query/acc.cgi?acc=GSE187022>, GSM5667182, GSM5667183, GSM5667184).

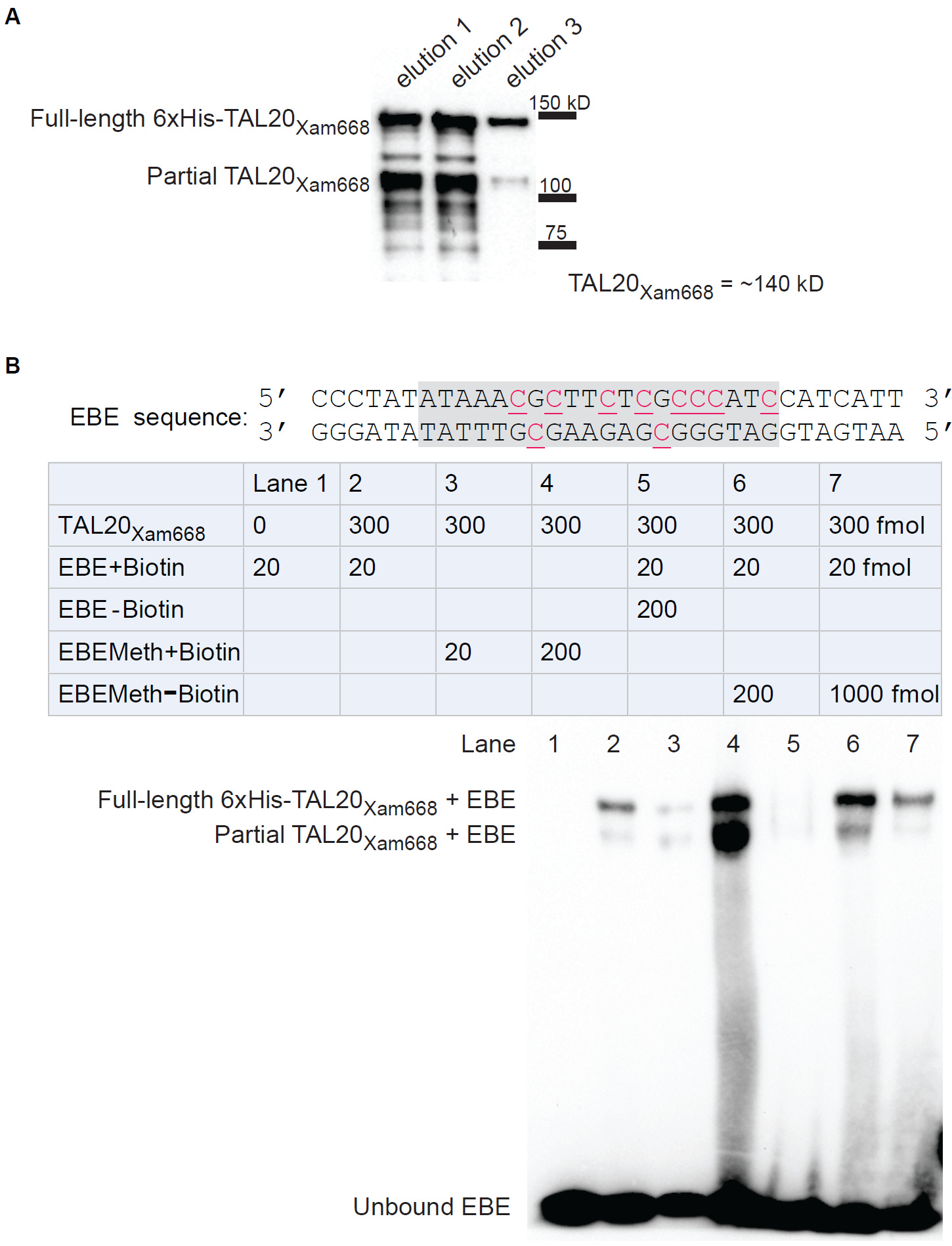

Supplementary Fig. 1 – Binding affinity of TAL20 to EBE is decreased *in vitro* in the presence of methylation.

(**A**) Western blot (anti-His) of elution samples resulting from TAL20 purification from *E. coli*. Three elutions were performed and are shown. Size standards are shown to the right (kD). (**B**) Electrophoretic mobility shift assay (EMSA) between the EBE and His-purified TAL20 (elution 3, panel A). The binding reaction components are given in the table with amounts indicated. The DNA sequence used in the binding reactions is shown above the table with the EBE sequence shaded in grey and methylated cytosines indicated in red and underlined. Each reaction corresponds to a lane in the blot shown below. The lower band from the purification, presumed to be partial TAL20 peptide or some other 6xHis-containing product, appears in the EMSA as well, and are labeled as such to the left of the image. This experiment was performed twice with similar results.

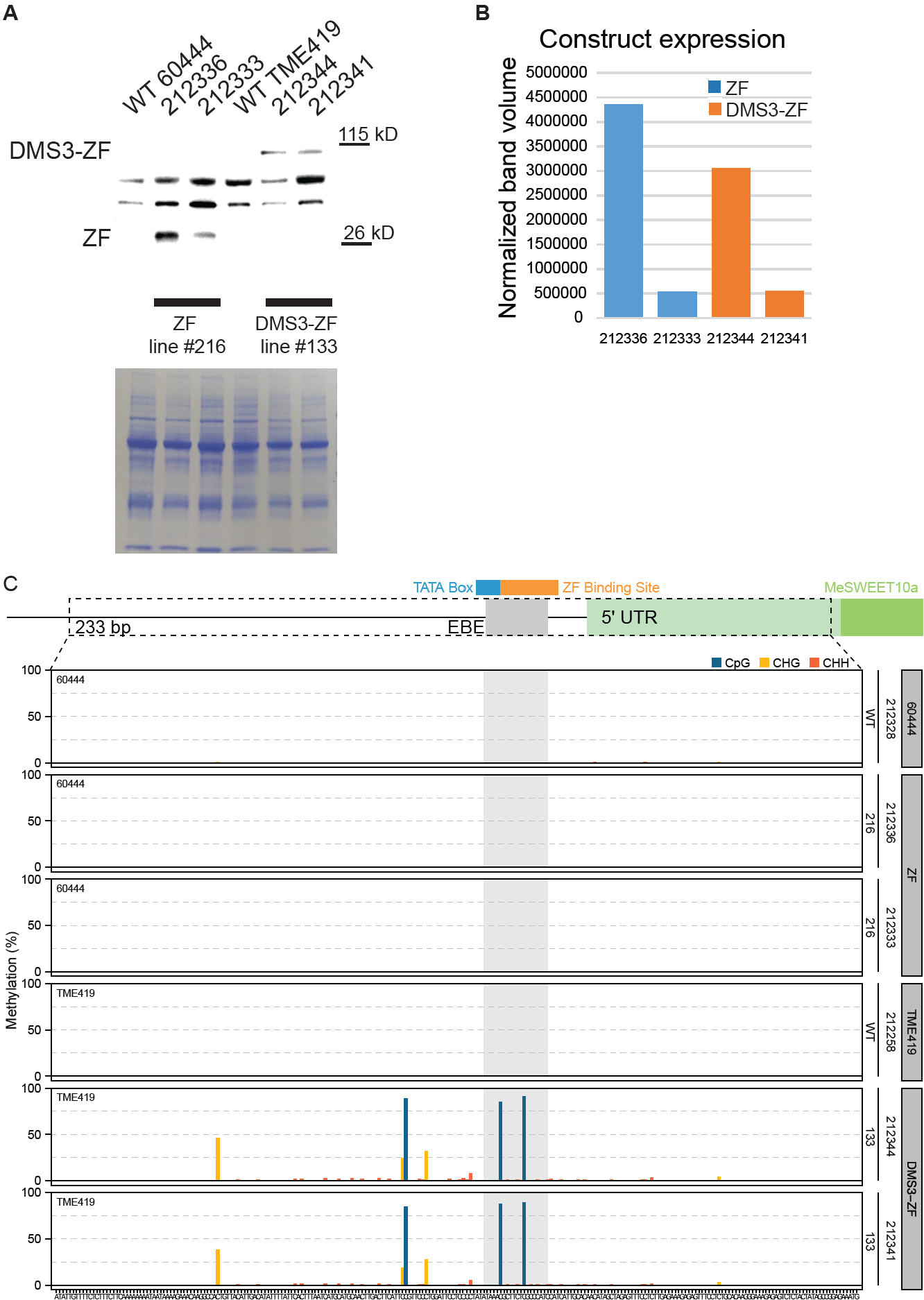

Supplementary Fig. 2 – Protein expression and methylation quantification for individuals used in an individual experiment (qPCR1).

(**A**) Expression of transgenes in individual plants from two independent DMS3-expressing transgenic lines (133 and 204) as well as a ZF-only negative control line (216). Top: Western blots (anti-FLAG) showing expression of the ZF (ZF-3xFLAG) protein with and without DMS3. Each individual plant is identified by a number (above each lane) and the necessary wild-type (WT) controls are also included (60444 for ZF and TME 419 for DMS3-ZF lines). Relevant size standards are shown to the right (kD). Bottom: Coomassie Brilliant Blue-stained Rubisco, loading control. (**B**) Quantification of the intensity of the bands, expressed as adjusted band volume (relative to Rubisco) according to Image Lab™ software (Bio-Rad). (**C**) PCR bisulfite sequencing (ampBS-seq) results from all samples shown in **A**. Top: Graphical depiction of *MeSWEET10a* promoter region assessed for methylation. The EBE (grey) which overlaps a presumed TATA box (blue), is indicated. The site that the ZF was engineered to bind is shown in orange. The predicted 5’ UTR and *MeSWEET10a* transcriptional start site are shown in green. The area within the dotted lined box (233 bp) was subjected to ampBS-seq. Bottom: CpG, CHG, and CHH DNA methylation levels (percent, y-axis) of the *MeSWEET10a* promoter (EBE, grey) measured by ampBS-seq with and without DMS3-ZF. Background of tissue for each plot is indicated to the right.

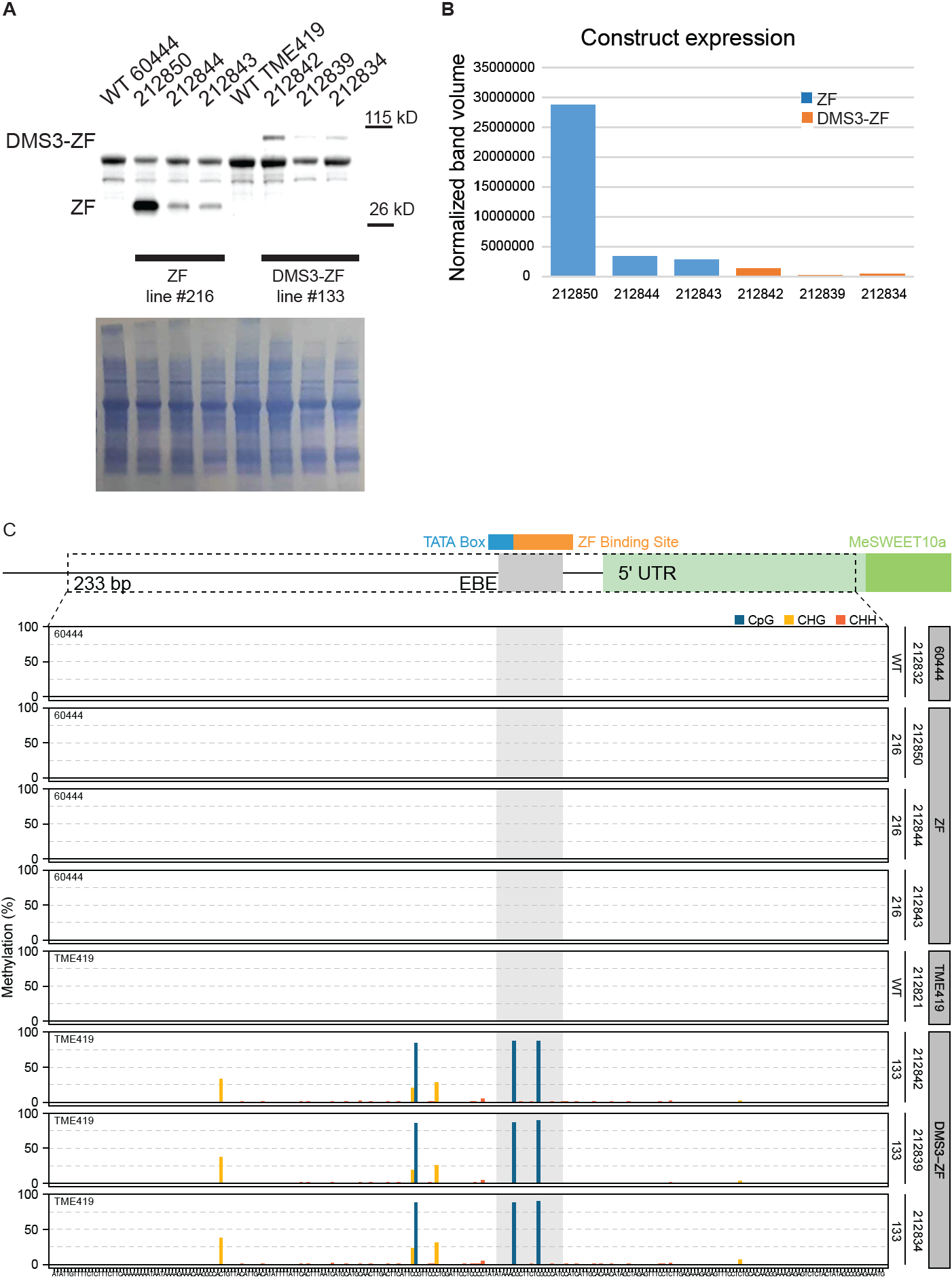

Supplementary Fig. 3 – Protein expression and methylation quantification for individuals used in an individual experiment (qPCR2).

(**A**) Expression of transgenes in individual plants from two independent DMS3-expressing transgenic lines (133 and 204) as well as a ZF-only negative control line (216). Top: Western blots (anti-FLAG) showing expression of the ZF (ZF-3xFLAG) protein with and without DMS3. Each individual plant is identified by a number (above each lane) and the necessary wild-type (WT) controls are also included (60444 for ZF and TME 419 for DMS3-ZF lines). Relevant size standards are shown to the right (kD). Bottom: Coomassie Brilliant Blue-stained Rubisco, loading control. (**B**) Quantification of the intensity of the bands, expressed as adjusted band volume (relative to Rubisco) according to Image Lab™ software (Bio-Rad). (**C**) PCR bisulfite sequencing (ampBS-seq) results from all samples shown in **A**. Top: Graphical depiction of *MeSWEET10a* promoter region assessed for methylation. The EBE (grey) which overlaps a presumed TATA box (blue), is indicated. The site that the ZF was engineered to bind is shown in orange. The predicted 5’ UTR and *MeSWEET10a* transcriptional start site are shown in green. The area within the dotted lined box (233 bp) was subjected to ampBS-seq. Bottom: CpG, CHG, and CHH DNA methylation levels (percent, y-axis) of the *MeSWEET10a* promoter (EBE, grey) measured by ampBS-seq with and without DMS3-ZF. Background of tissue for each plot is indicated to the right.

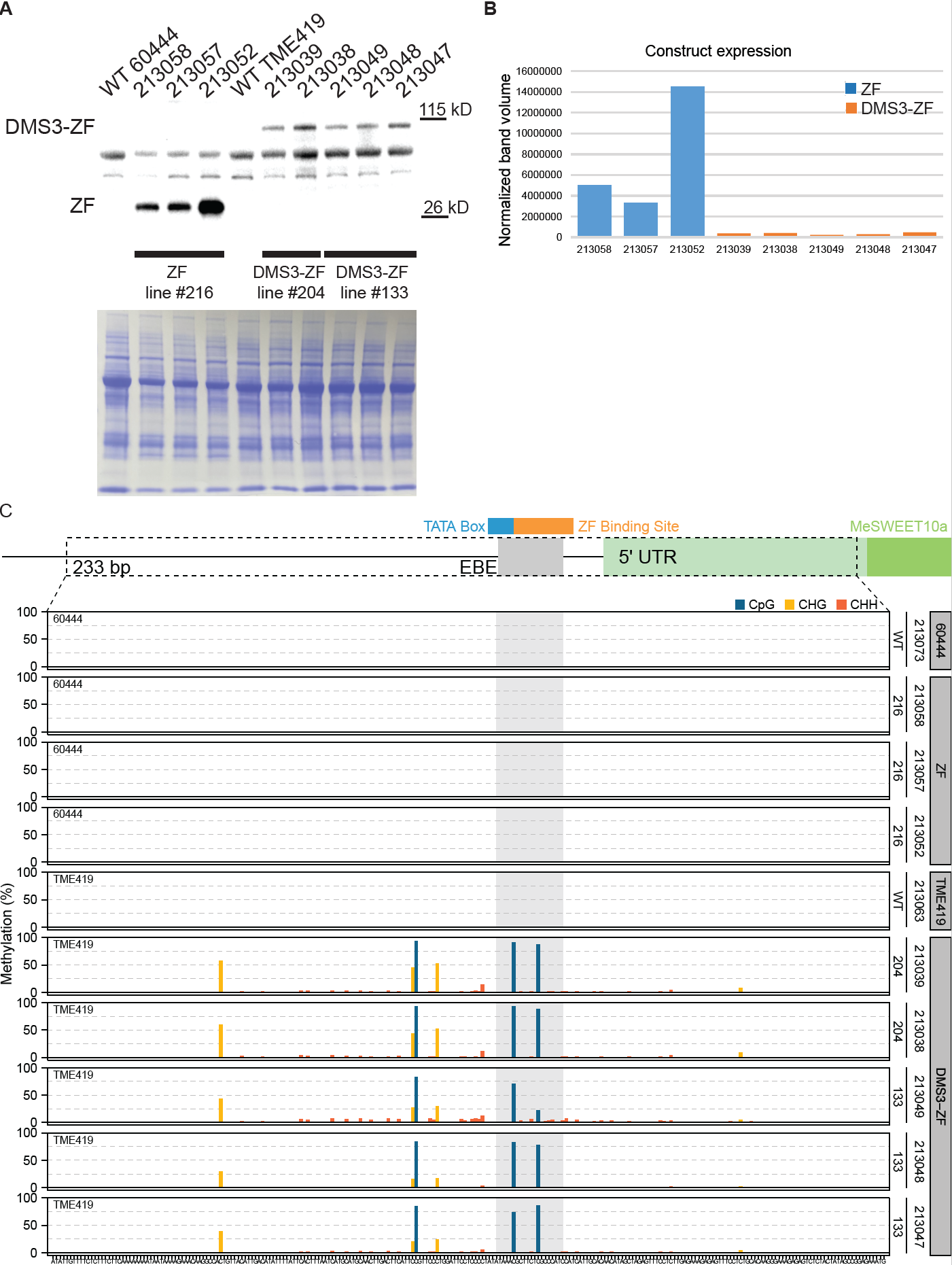

Supplementary Fig. 4 – Protein expression and methylation quantification for individuals used in an individual experiment (bacterial growth, water-soaking).

(**A**) Expression of transgenes in individual plants from two independent DMS3-expressing transgenic lines (133 and 204) as well as a ZF-only negative control line (216). Top: Western blots (anti-FLAG) showing expression of the ZF (ZF-3xFLAG) protein with and without DMS3. Each individual plant is identified by a number (above each lane) and the necessary wild-type (WT) controls are also included (60444 for ZF and TME 419 for DMS3-ZF lines). Relevant size standards are shown to the right (kD). Bottom: Coomassie Brilliant Blue-stained Rubisco, loading control. (**B**) Quantification of the intensity of the bands, expressed as adjusted band volume (relative to Rubisco) according to Image Lab™ software (Bio-Rad). (**C**) PCR bisulfite sequencing (ampBS-seq) results from all samples shown in **A**. Top: Graphical depiction of *MeSWEET10a* promoter region assessed for methylation. The EBE (grey) which overlaps a presumed TATA box (blue), is indicated. The site that the ZF was engineered to bind is shown in orange. The predicted 5’ UTR and *MeSWEET10a* transcriptional start site are shown in green. The area within the dotted lined box (233 bp) was subjected to ampBS-seq. Bottom: CpG, CHG, and CHH DNA methylation levels (percent, y-axis) of the *MeSWEET10a* promoter (EBE, grey) measured by ampBS-seq with and without DMS3-ZF. Background of tissue for each plot is indicated to the right.

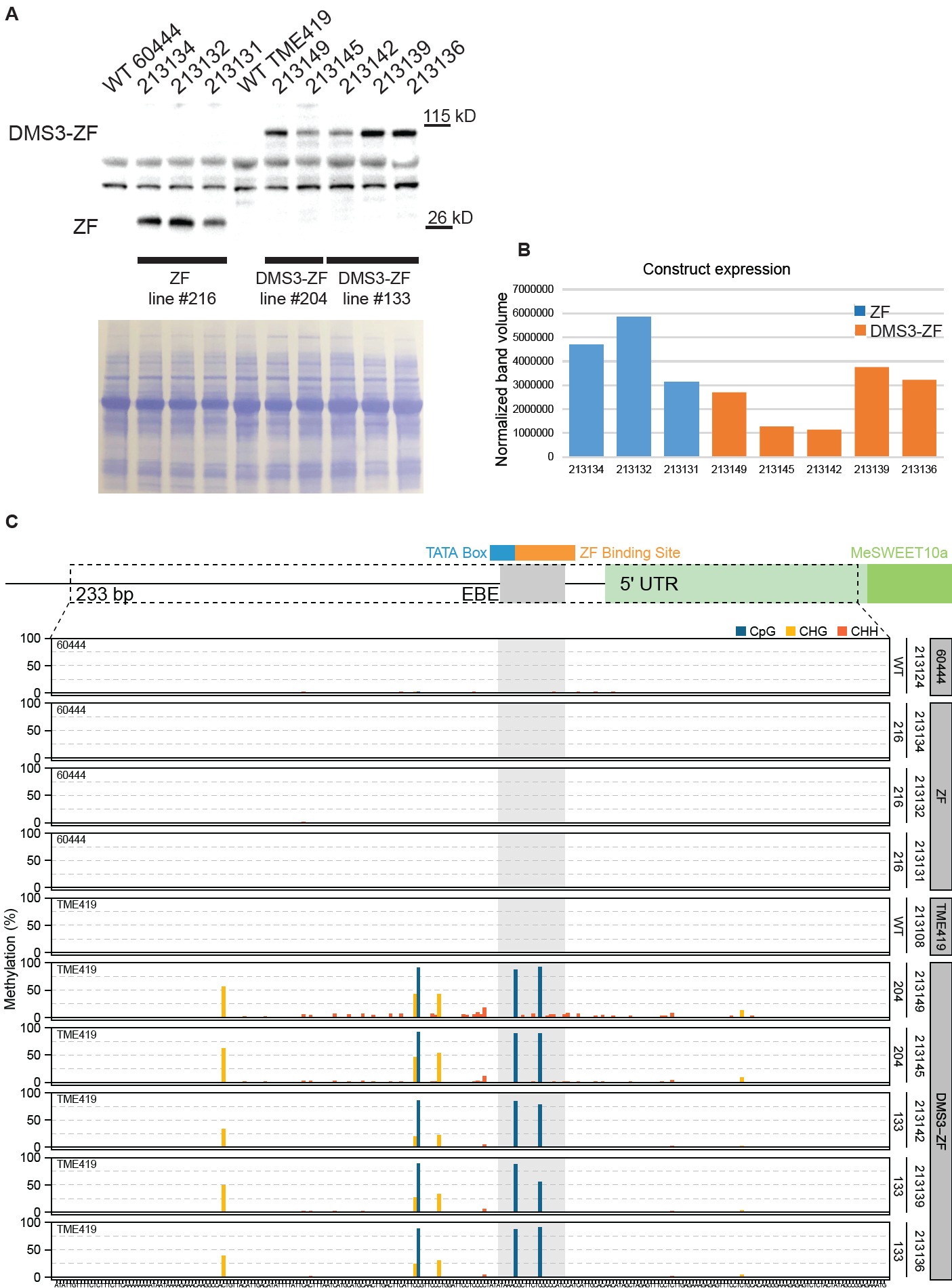

Supplementary Fig. 5 – Protein expression and methylation quantification for individuals used in an individual experiment (qPCR3).

(**A**) Expression of transgenes in individual plants from two independent DMS3-expressing transgenic lines (133 and 204) as well as a ZF-only negative control line (216). Top: Western blots (anti-FLAG) showing expression of the ZF (ZF-3xFLAG) protein with and without DMS3. Each individual plant is identified by a number (above each lane) and the necessary wild-type (WT) controls are also included (60444 for ZF and TME 419 for DMS3-ZF lines). Relevant size standards are shown to the right (kD). Bottom: Coomassie Brilliant Blue-stained Rubisco, loading control. (**B**) Quantification of the intensity of the bands, expressed as adjusted band volume (relative to Rubisco) according to Image Lab™ software (Bio-Rad). (**C**) PCR bisulfite sequencing (ampBS-seq) results from all samples shown in **A**. Top: Graphical depiction of *MeSWEET10a* promoter region assessed for methylation. The EBE (grey) which overlaps a presumed TATA box (blue), is indicated. The site that the ZF was engineered to bind is shown in orange. The predicted 5’ UTR and *MeSWEET10a* transcriptional start site are shown in green. The area within the dotted lined box (233 bp) was subjected to ampBS-seq. Bottom: CpG, CHG, and CHH DNA methylation levels (percent, y-axis) of the *MeSWEET10a* promoter (EBE, grey) measured by ampBS-seq with and without DMS3-ZF. Background of tissue for each plot is indicated to the right.

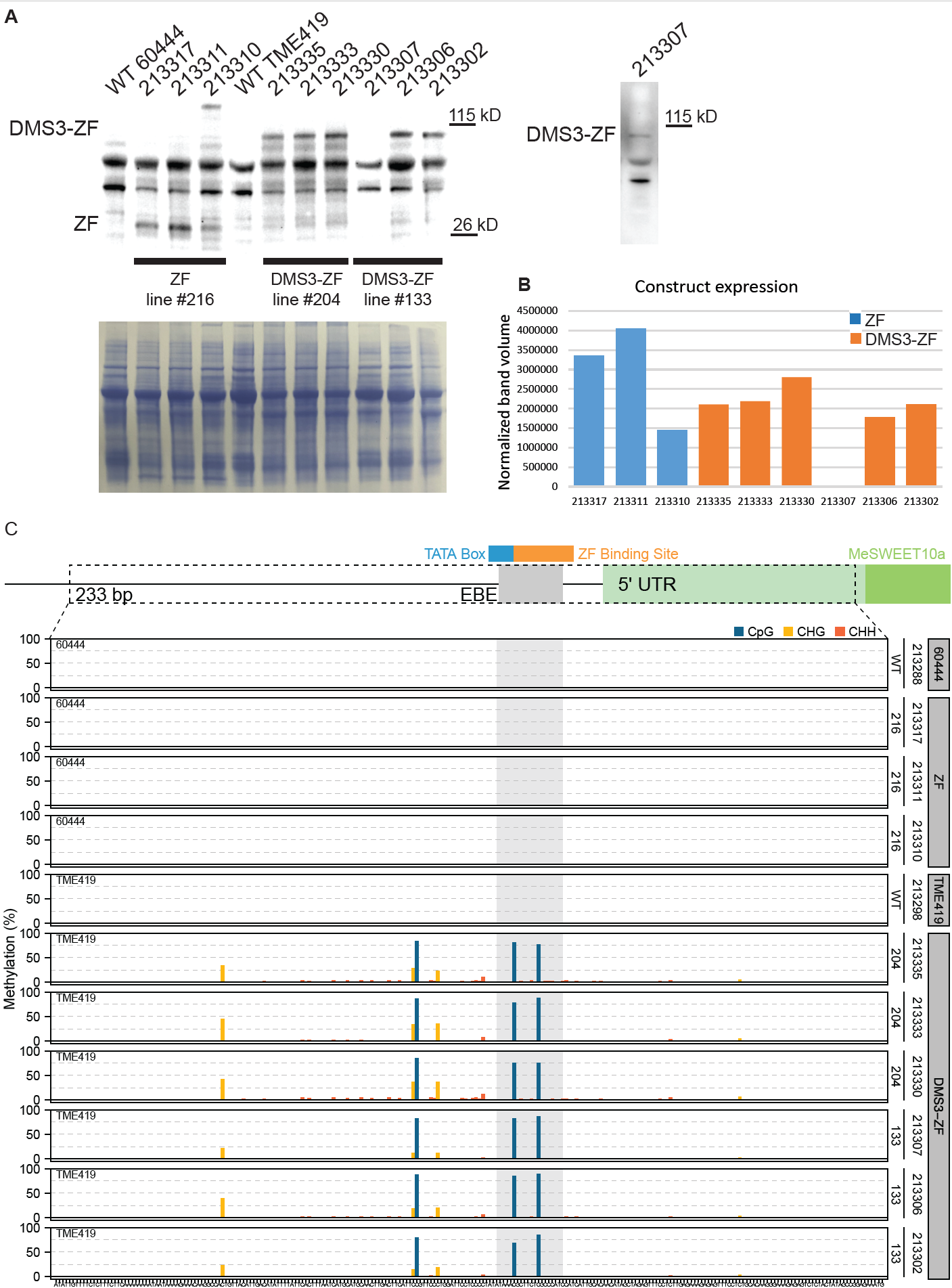

Supplementary Fig. 6 – Protein expression and methylation quantification for individuals used as an individual experiment set (bacterial growth, qPCR4, water-soaking).

(**A**) Expression of transgenes in individual plants from two independent DMS3-expressing transgenic lines (133 and 204) as well as a ZF-only negative control line (216). Top: Western blots (anti-FLAG) showing expression of the ZF (ZF-3xFLAG) protein with and without DMS3. Each individual plant is identified by a number (above each lane) and the necessary wild-type (WT) controls are also included (60444 for ZF and TME 419 for DMS3-ZF lines). Relevant size standards are shown to the right (kD). Bottom: Coomassie Brilliant Blue-stained Rubisco, loading control. Right: Example replicate blot showing expression of DMS3-ZF in individual 213307. No expression of DMS3-ZF was detected in individual 213307 in the blot used for quantification. Additional repeats of western blot analysis on that sample showed evidence of detectable expression of DMS3-ZF. (**B**) Quantification of the intensity of the bands shown in **A**, expressed as adjusted band volume (relative to Rubisco) according to Image Lab™ software (Bio-Rad). (**C**) PCR bisulfite sequencing (ampBS-seq) results from all samples shown in **A**. Top: Graphical depiction of *MeSWEET10a* promoter region assessed for methylation. The EBE (grey) which overlaps a presumed TATA box (blue), is indicated. The site that the ZF was engineered to bind is shown in orange. The predicted 5’ UTR and *MeSWEET10a* transcriptional start site are shown in green. The area within the dotted lined box (233 bp) was subjected to ampBS-seq. Bottom: CpG, CHG, and CHH DNA methylation levels (percent, y-axis) of the *MeSWEET10a* promoter (EBE, grey) measured by ampBS-seq with and without DMS3-ZF. Background of tissue for each plot is indicated to the right.

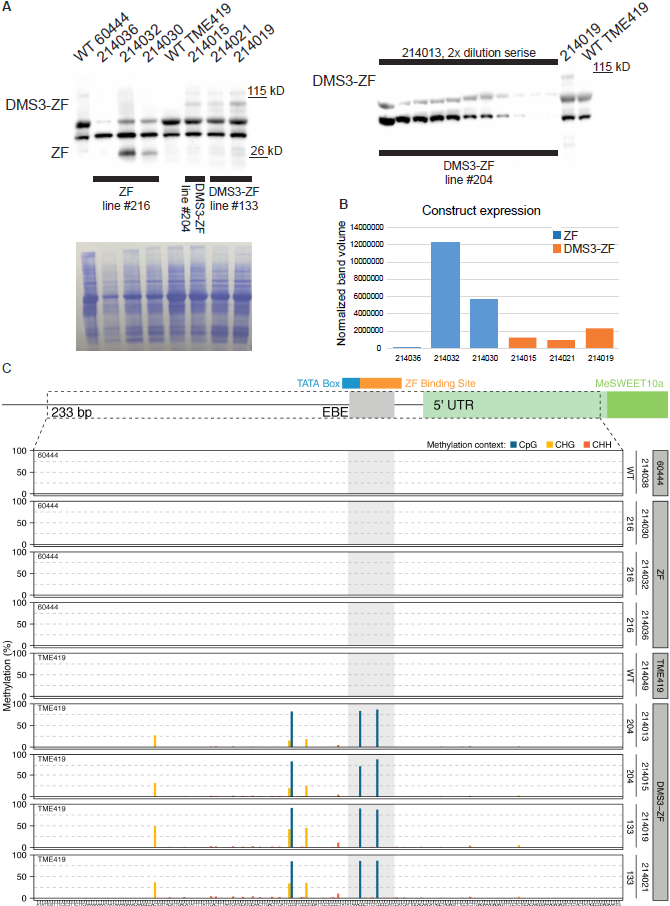

Supplementary Fig. 7 – Protein expression and methylation quantification for individuals used in an individual experiment (bacterial growth, water-soaking).

(**A**) Expression of transgenes in individual plants from two independent DMS3-expressing transgenic lines (133 and 204) as well as a ZF-only negative control line (216). Top: Western blots (anti-FLAG) showing expression of the ZF (ZF-3xFLAG) protein with and without DMS3. Each individual plant is identified by a number (above each lane) and the necessary wild-type (WT) controls are also included (60444 for ZF and TME 419 for DMS3-ZF lines). Relevant size standards are shown to the right (kD). Bottom: Coomassie Brilliant Blue-stained Rubisco, loading control. Right: Example dilution series blot showing the lack detectable expression of DMS3-ZF in an individual (214013) included in a water-soaking experiment. (**B**) Quantification of the intensity of the bands shown in A, expressed as adjusted band volume (relative to Rubisco) according to Image Lab™ software (Bio-Rad). (**C**) PCR bisulfite sequencing (ampBS-seq) results from all samples shown in **A**. Top: Graphical depiction of *MeSWEET10a* promoter region assessed for methylation. The EBE (grey) which overlaps a presumed TATA box (blue), is indicated. The site that the ZF was engineered to bind is shown in orange. The predicted 5’ UTR and *MeSWEET10a* transcriptional start site are shown in green. The area within the dotted lined box (233 bp) was subjected to ampBS-seq. Bottom: CpG, CHG, and CHH DNA methylation levels (percent, y-axis) of the *MeSWEET10a* promoter (EBE, grey) measured by ampBS-seq with and without DMS3-ZF. Background of tissue for each plot is indicated to the right. Note: the observed methylation in individual 214013 is indistinguishable from other individuals, despite undetectable expression of the DMS3-ZF construct.

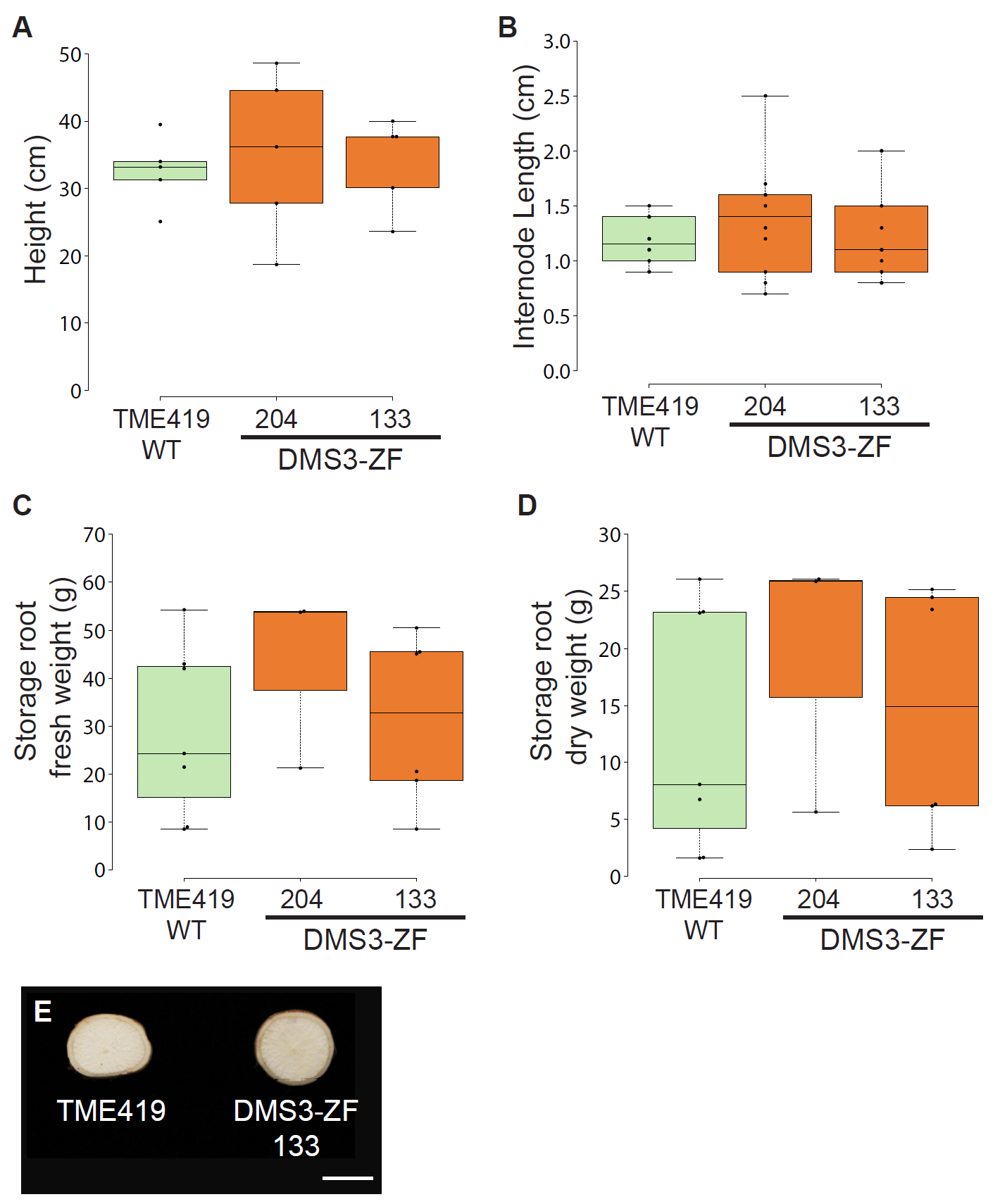

**Supplementary Fig. 8 – Developmental characteristics are indistinguishable between wild-type and methylated transgenic lines.**

**(A)** Height and **(B)** internode length of individual wild-type (WT, green) and DMS3-ZF (orange), field-grown plants after approximately 6 months in soil. Internode measurements for each plant were taken on the stem just above and below the woody transition. Biological replicate values are indicated by dots (n = 5 per background). Horizontal black line within boxes indicates the value of the median while the box limits indicate the 25th and 75th percentiles as determined by R software; whiskers extend 1.5 times the interquartile range (1.5xIQR) from the 25th and 75th percentiles. **(C)** Fresh and **(D)** dry weight of storage roots induced in a greenhouse setting. Dry weight of cassava storage roots consists of approximately 80% starch. No significant differences were detected for any trait measured (1-way ANOVA, *p* >0.05). Biological replicates (black dots) included in each background (x-axis) are as follows: n = 7, 3, 6. **(E)** Representative image of fresh storage root cross-section measured in panels C and D. Scale bar = 1.0 cm.

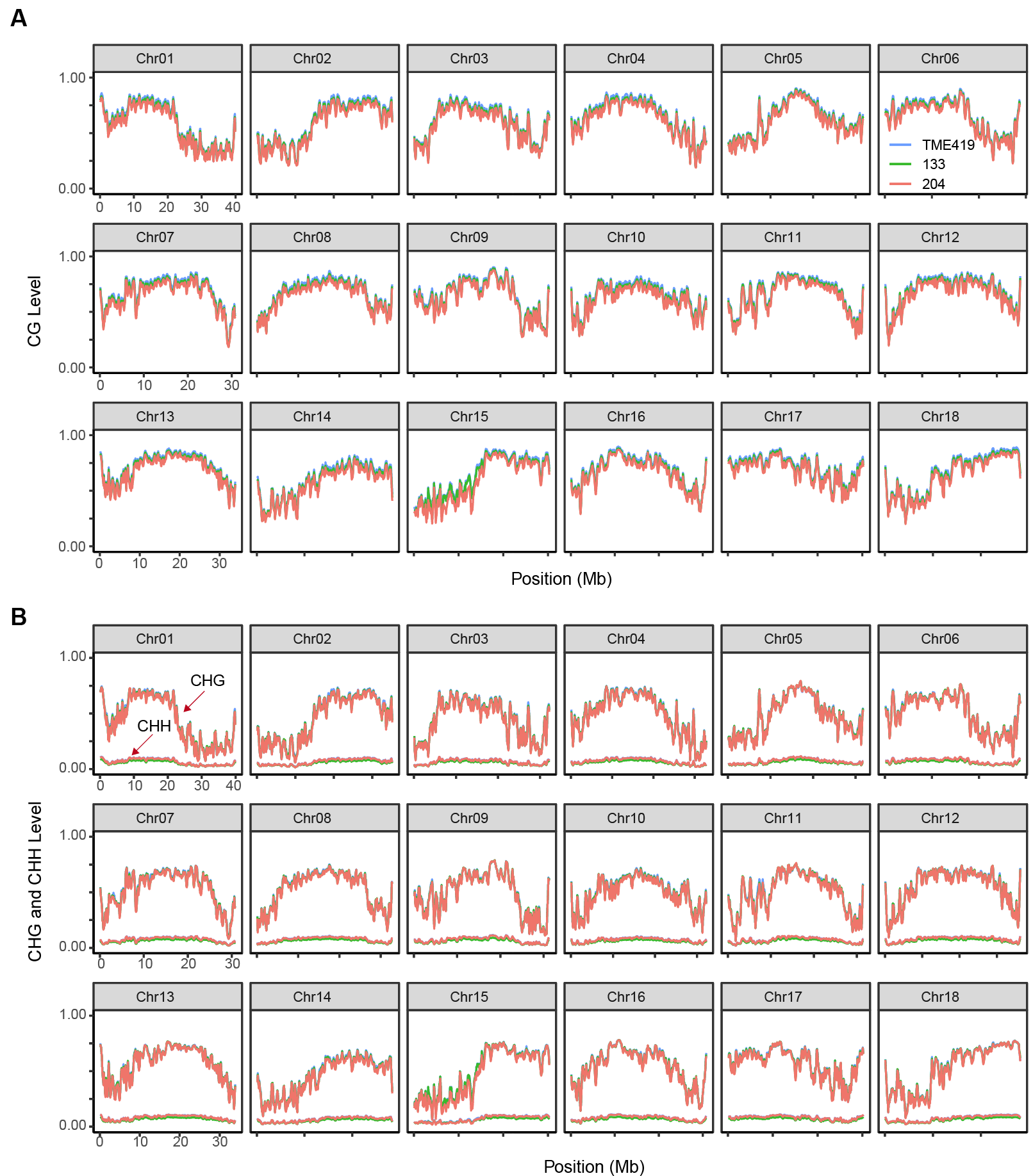

**Supplementary Fig. 9 – Genome-wide DNA methylation of chromosomes.**

Genome-wide distribution of **(A)** CpG, **(B)** CHG, and CHH methylation over 18 Cassava chromosomes in wild-type TME419 and two independent DMS3-ZF lines (133 and 204). Methylation levels were calculated by splitting chromosomal sequences into 100 kb bins.

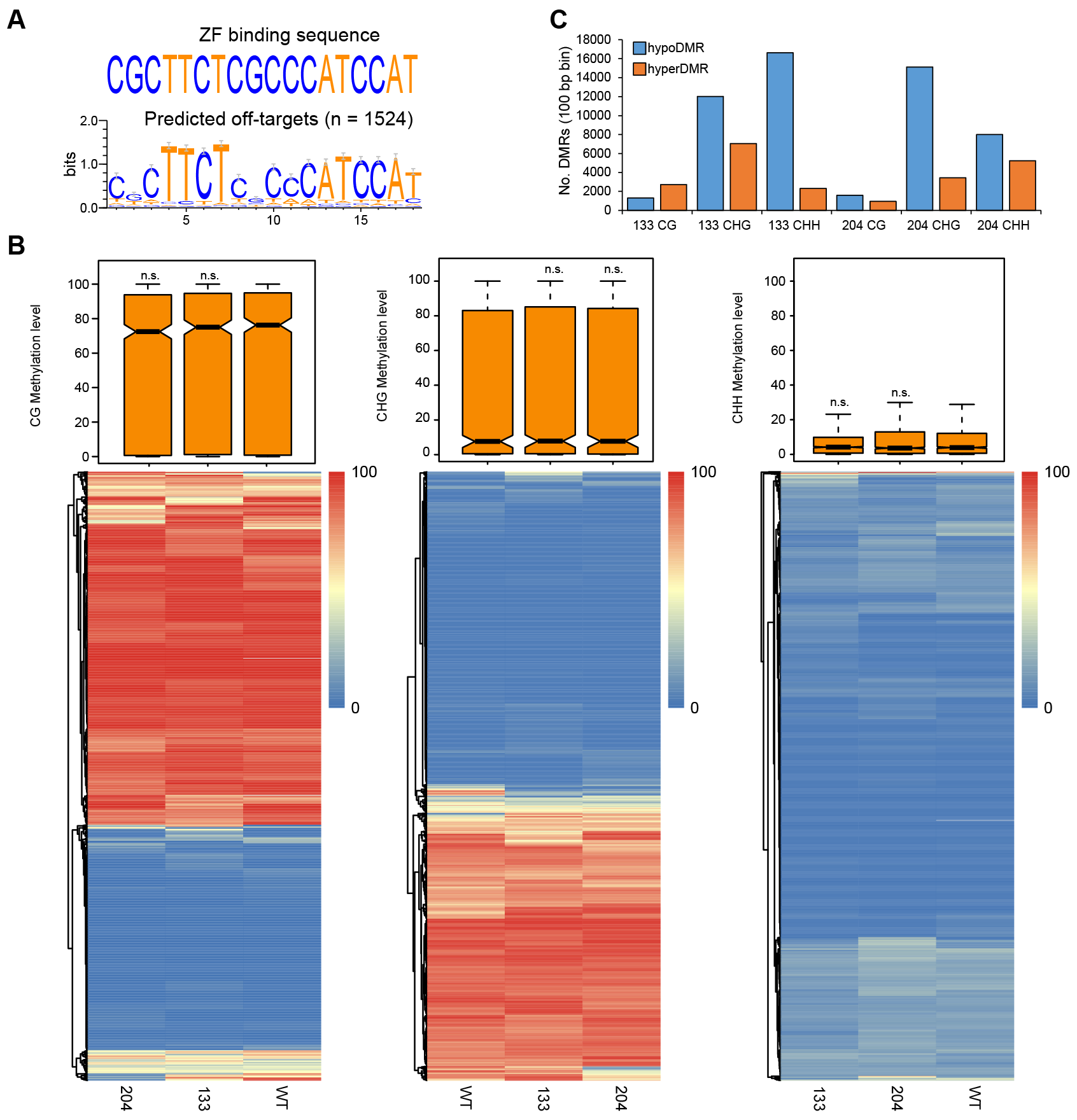

**Supplementary Fig. 10 – Off-target and differentially methylated region (DMR) analysis. (A)** Logo representation of genome-wide prediction of potential ZF off-target binding sites. The position within the sequence is plotted on the x-axis, and y-axis shows the number of bits of information. The height of each nucleotide at each position within the target sequence represents the calculated level of sequence conservation. Error bars are Bayesian 95% confidence intervals. **(B)** Box plot and heatmap showing CG, CHG, and CHH methylation percentage (WGBS data, y-axis) of ZF off-targets in wild-type TME419 (WT) and two DMS3-ZF-expressing lines (133 and 204). The horizontal black line within boxes indicates the median and the notch represents the 95% confidence interval of the median. The box limits indicate the 25th and 75th percentiles as determined by R software; whiskers extend 1.5 times the interquartile range (1.5xIQR) from the 25th and 75th percentiles. n = 1524 potential off-target sites were examined between TME419 (WT) and two DMS3-ZF-expressing lines (133 and 204). Insignificant comparisons (two-sided Student’s *t*-test without multiple comparisons adjustments) are labeled (n.s.). The heatmap shows CG, CHG, and CHH methylation percentage of potential ZF off-target binding sites in wild-type TME419 (WT) and two DMS3-ZF-expressing lines (133 and 204). **(C)** Bar chart showing the number of hypo- (blue) or hyper- (orange) CG, CHG, and CHH differentially methylated regions (DMRs) in two DMS3-ZF plant lines (133 and 204). All DMRs are 100 bp bins (y-axis) with CG variation > 0.4, CHG variation > 0.2, and CHH variation > 0.1.

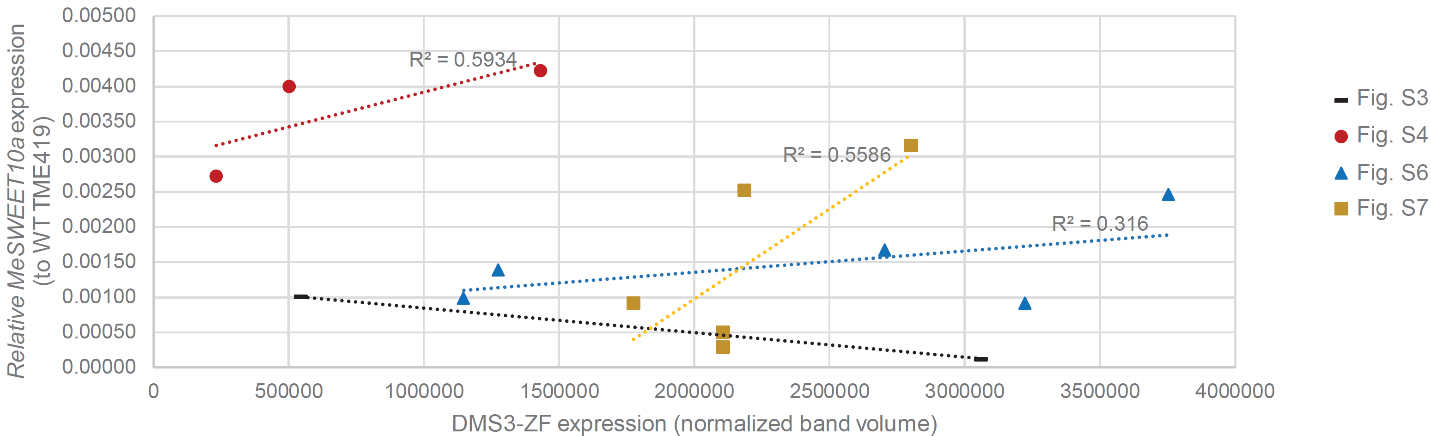
 **Supplementary Fig. 11 – Higher DMS3-ZF expression does not correlate with reduced *MeSWEET10a* induction.**

All *MeSWEET10a* expression values (y-axis) from Fig. 4 (*Xam*-treated leaves) plotted against the DMS3-ZF expression level quantified in **Supplementary Figs. 3, 4, 6,** and **7** (qPCR experiments 1-4). The legend (right) identifies each experiment. An individual plant with undetectable expression from Fig. S7 (214013) was not included in this analysis. R^2^ values suggest a lack of correlation for each experimental set, where applicable.

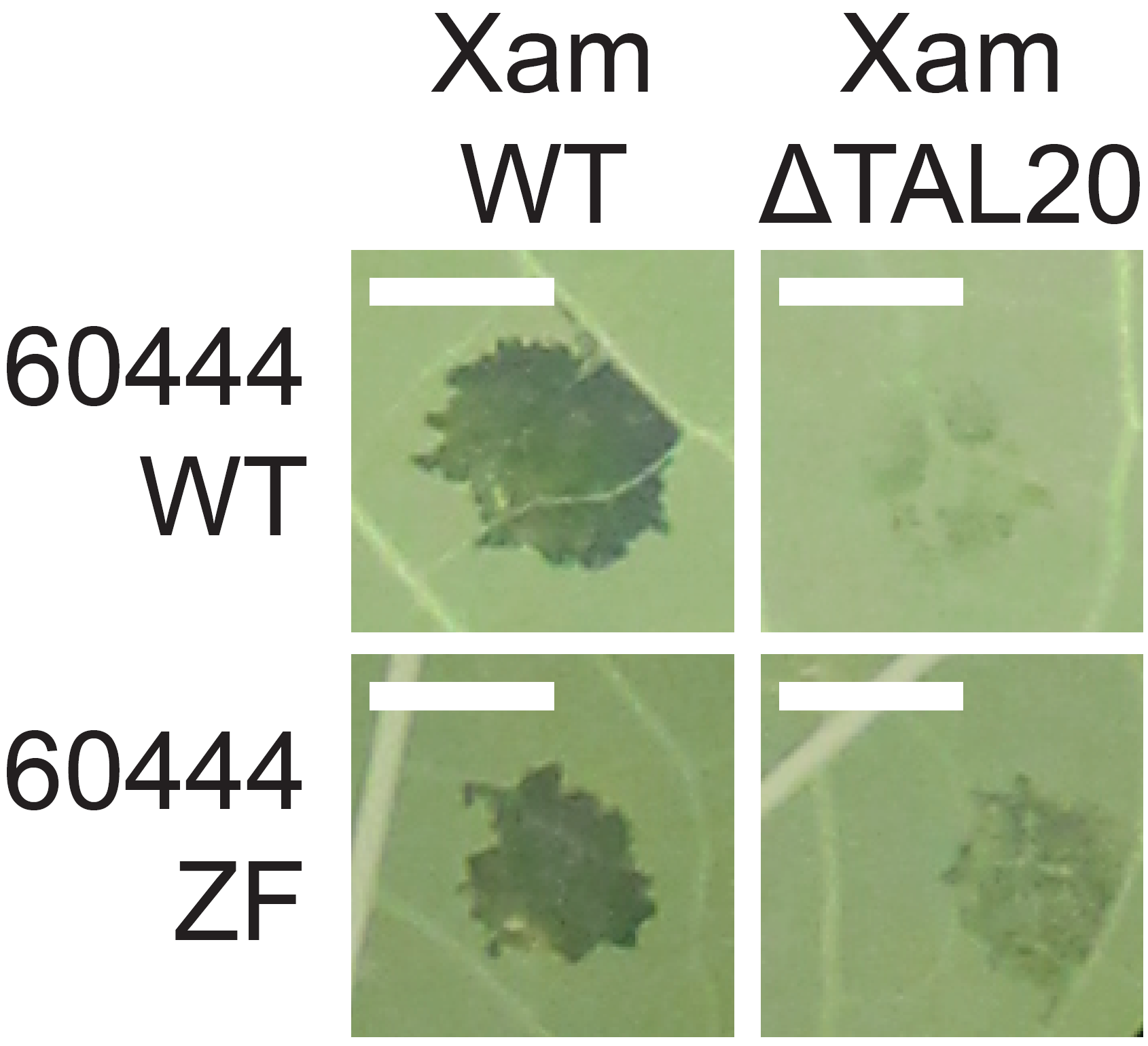

Supplementary Fig. 12 – Phenotype of additional *Xam*-infiltrated control plants.

Representative images of water-soaking phenotype of leaves from 60444 wild-type (WT) and ZF negative control plants. Images were taken 4-days post-infection with either *Xam668* (XamWT) or a *Xam668* TAL20 deletion mutant (XamΔTAL20) and originate from the six independent experiments presented in the manuscript. Scale bar = 0.5 cm.

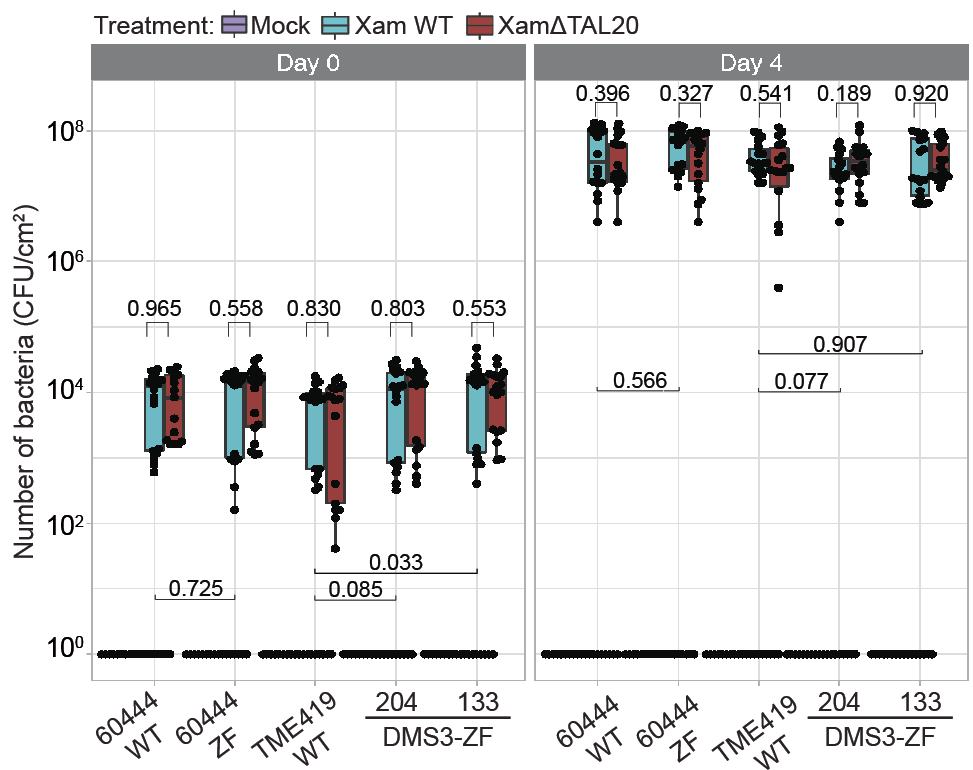

**Supplementary Fig. 13 – Bacterial growth is unaffected by methylation at the EBE.**

Bacterial populations in leaves measured at day-0 (left) and day-4 (right) post-infiltration with Xam (treatments listed above plot). Mean colony forming units (CFU/cm^2^, y-axis) are plotted per background tested (x-axis). Individual data points are represented as black dots analyzed across 3 independent experiments. For each background from left to right, Day 0 Mock, Xam WT, and XamΔTAL20 n = (18, 18, 15), n = (16,18,18), n = (18,18,17), n = (18,18,18), and n = (18,18,18). For each background from left to right, Day 4 Mock, Xam WT, and XamΔTAL20 n = (18,18,18), n = (18,18,18), n = (18,18,18), n = (18,16,18), and n = (16,18,18). Dots outside whiskers represent outliers. The horizontal line within the box represents the median sample value. The ends of the boxes represent the 3rd (Q3) and 1st (Q1) quartiles. The whiskers show values that are 1.5 times interquartile range (1.5xIQR) above and below Q1 and Q3. Results of statistical analyses (*p*-values, Student’s *t*-test) comparing the difference between treatments within each background (black text, above boxes) and the difference between Xam WT growth across different backgrounds (below boxes) are shown.

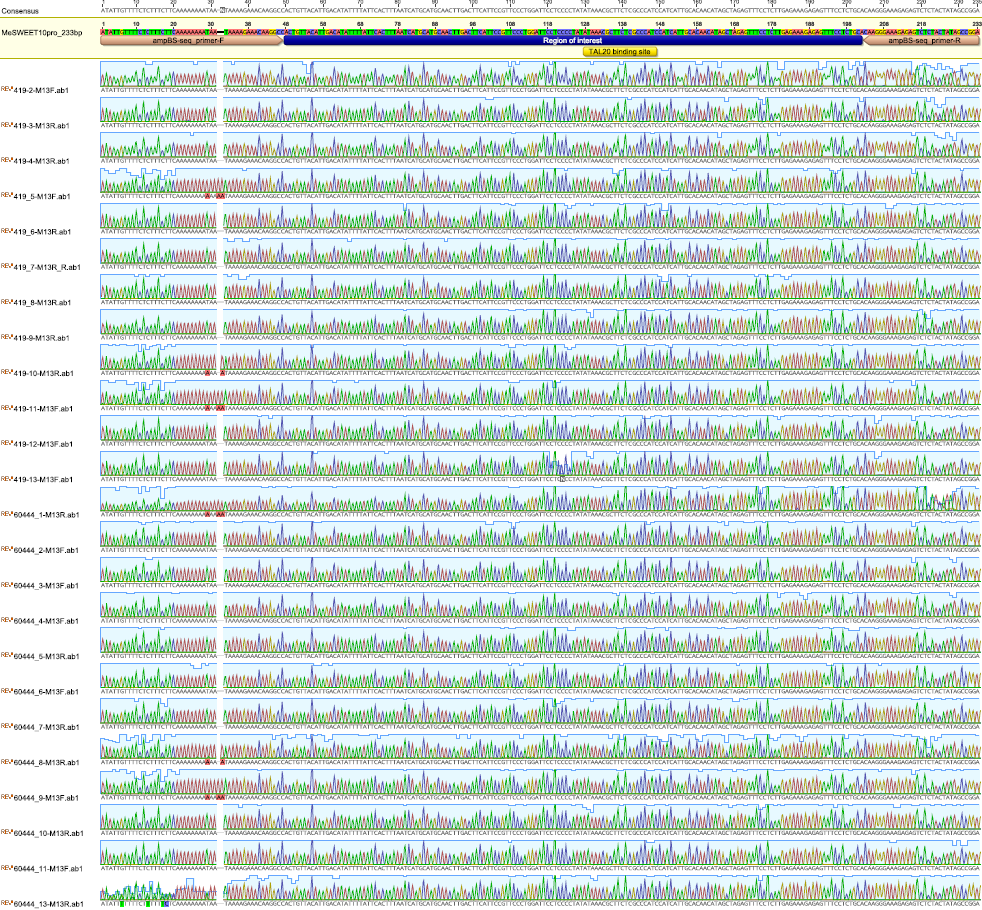

Supplementary Fig. 14 – *MeSWEET10a* promoter region of interest is identical in 60444 and TME419 WT backgrounds.

Sanger sequencing verifying the sequence within the WT backgrounds used in this study (60444 and TME419). Traces from 12 individual clones from each background are shown. No sequence variation was found within in region of interest.

.

**Supplementary Table 1 – Library information of WGBS coverage.**

| **Sample** | **Reads count** | **Library** | **Coverage*** | **Conversion rate**** | **GEO ID** |
| --- | --- | --- | --- | --- | --- |
| WT | 573172769 | PE100 | 159.00 | 98.24% | GSM5667182 |
| ZF133 | 403083073 | PE100 | 111.82 | 97.42% | GSM5667183 |
| ZF204 | 467616824 | PE100 | 129.72 | 98.01% | GSM5667184 |

* Estimated with genome size: 720,958,040 bp. ** Estimated with chloroplast genome.

Supplementary Table 2 - Overlapping of DMRs with ZF off-targets and random shuffling control.

| **Line_context_type** | **Overlap with predicted off-targets (n =1524)** | **Overlap with random control (n =1524)** |
| --- | --- | --- |
| ZF133_CG_hyperDMR | 1 | 1 |
| ZF133_CG_hypoDMR | 0 | 0 |
| ZF133_CHG_hyperDMR | 5 | 6 |
| ZF133_CHG_hypoDMR | 5 | 7 |
| ZF133_CHH_hyperDMR | 1 | 2 |
| ZF133_CHH_hypoDMR | 14 | 9 |
| ZF204_CG_hyperDMR | 0 | 0 |
| ZF204_CG_hypoDMR | 3 | 1 |
| ZF204_CHG_hyperDMR | 2 | 3 |
| ZF204_CHG_hypoDMR | 10 | 7 |
| ZF204_CHH_hyperDMR | 6 | 3 |
| ZF204_CHH_hypoDMR | 7 | 9 |

Supplementary Table 3 - Primers used to generate data presented in this manuscript.

| **Primer Number** | **Primer Name** | **Sequence (5'-3')** | **Description** |
| --- | --- | --- | --- |
| 107 | Manes.09G039900_Me PP2A-4 NTv2_For | AGGCTCACACTTTCATCCAGTTTGAG | RT-qPCR |
| 108 | Manes.09G039900_Me PP2A-4 NTv2_Rev | ACCTGAGCGTAAAGCAGGGAAG | RT-qPCR |
| 109 | GTPb (Manes.09G086600)_For | CCTCAAAGGCTGAGCCACAGA | RT-qPCR |
| 110 | GTPb (Manes.09G086600)_Rev | GGGAGAAACAATACAGGCACCAATCAC | RT-qPCR |
| 348 | 10a_qPCR-F3 | GCGGTGATGTGGTTCTTC | RT-qPCR |
| 349 | 10a_qPCR-R3 | CGATGTGCTCGGACAATTC | RT-qPCR |
| 117 | Manes.15G048700_qPCR-F3 | CCTGGTTGATGCTGTCATGGG | RT-qPCR |
| 119 | Manes.15G048700_qPCR-R4 | GGTGGGATTTGCACTTCCACC | RT-qPCR |
| 120 | JP16889 | ATATTGTTTTTTTTTTTTTTAAAAAAAATAATAAAAGAAATAAGGTT | ampBS-seq |
| 121 | JP16890 | TCCRACTATAATAAAAACTCTCTTTCCCTTA | ampBS-seq |

Each primer has a number and a name (first two columns). Column 3: the sequence of the primer (5’ to 3’ direction). Column 4: How primer was used.
